# Hybridization load introduced by Alpine ibex hybrid swarms

**DOI:** 10.1101/2024.06.24.599926

**Authors:** F. Gözde Çilingir, Fabio Landuzzi, Alice Brambilla, Debora Charrance, Federica Furia, Sara Trova, Alberto Peracino, Glauco Camenisch, Dominique Waldvogel, Jo Howard-McCombe, Yeraldin Chiquinquira Castillo De Spelorzi, Edoardo Henzen, Andrea Bernagozzi, Alessandro Coppe, Jean Marc Christille, Manuela Vecchi, Diego Vozzi, Andrea Cavalli, Bruno Bassano, Stefano Gustincich, Daniel Croll, Luca Pandolfini, Christine Grossen

## Abstract

Species conservation efforts can be threatened by deleterious mutation accumulation following population contractions. In addition to *de novo* mutations, a significant source of genetic load could be deleterious variants introduced into a population through hybridization. Hence, even successfully restored species may face deleterious mutation swamping due to hybridization with an abundant and closely related species. The outcomes of such hybridization events are poorly understood given the complex interplay of introduced adaptive and maladaptive variation. Here, we analyze this potential risk for Alpine ibex (*Capra ibex*), a flagship species of large mammal restoration in the Alps. Near-extinction two centuries ago resulted in exceptionally low genome-wide diversity and increased inbreeding, which facilitated the purging of severe deleterious mutations but accumulation of less severe ones. We produced a highly contiguous chromosome-level genome assembly of the Alpine ibex capturing structural divergence from its closest domestic species, the domestic goat (*Capra hircus*) known to hybridize with Alpine ibex Genome sequencing of eight recent ibex-goat hybrids and backcrosses from two hybrid swarms in Northern Italy revealed highly diverse recombinants and an average of 30 masked, predicted loss-of-function (LOF) mutations per hybrid compared to 10 in non-hybrid Alpine ibex. This exposes Alpine ibex to further backcrosses, exposing their vulnerable gene pool to an influx of hybridization load. Individual-based genomic simulations suggest that such LOF load would return to pre-hybridization levels with a lag of over 100 generations after gene flow subsides. Hybridization could also disrupt local adaptation in the recipient species. Our work provides a direct estimate of hybridization load and, by this, informs on the complexity of managing endangered gene pools in the face of hybridization.

## Introduction

Human activity has profoundly affected species and populations through habitat degradation and over-exploitation leading to severe population reductions and extinctions^1^. A particular concern for species conservation is increased levels of inbreeding and the expression of deleterious mutations in small populations. Recent studies provided compelling evidence for changes in deleterious mutation load in natural populations (reviewed in ^2,3^). As predicted by theory, population bottlenecks were repeatedly shown to lead to the selective removal (purging) of highly deleterious (usually recessive) mutations, while less severely deleterious mutations can accumulate (*e.g.,* ^4–6^; reviewed in ^2^). Mutation load is defined as load resulting from newly introduced deleterious mutations^7^. Empirical assessments of fitness effects confirmed the detrimental impact of mutation load and inbreeding on both individual and population fitness^8–10^. However, the origin of deleterious mutations remains often unresolved. In addition to *de novo* mutations, a potentially significant source of mutation load could be deleterious variants introduced into a population through hybridization.

Hybridization in nature contributes to genetic variation and fosters adaptation^11,12^. Hybridization in bottlenecked species may introduce new adaptive genetic variation, compensating previously lost variation^13^. In addition, high levels of heterozygosity in hybrids can also initially mask deleterious mutations, thereby causing hybrid vigor (*i.e.*, heterosis) and mitigating inbreeding. Yet, this initial masking of deleterious mutations prevents the counter-selection of deleterious mutations in early generations. Deleterious mutations likely also played a significant role in hybridization events in animals such as fish^14,15^, plants including crops^16–18^, and trees^19^. These studies indicate that while hybridization can introduce beneficial alleles, linked deleterious variants can reduce fitness and create partial barriers to introgression. Hybridization also poses the risk of disrupting local adaptation through introgression of non-native alleles, in particular if the new species contact is caused by human activities^20–23^. Such a swamping of the native gene pool by hybridization is evident in Scottish wildcat and affects, among others, immunity-related functions^24^. The introduction of deleterious mutations is defined as hybridization load^13^. The interplay of adaptive and maladaptive variation introduced through hybridization is likely complex and largely lacks direct evidence from ongoing hybridization events.

To quantify load in genomes and to predict effects of introgression, high-quality genomic resources are essential^25^. The Alpine ibex (*Capra ibex*), a mountain ungulate endemic to the European Alps (Figure 1A, B), has experienced severe bottlenecks since the early 19th century and repeated hybridization events^26,27^. Conservation efforts have restored the species to its original distribution area, with a current population of about 55,000 individuals^28^. However, the past near extinction resulted in Alpine ibex having one of the lowest levels of genetic variation among mammals^5,29^. This reduced genetic diversity has led to high among-individual genetic similarity resulting in genomic signatures of inbreeding. Furthermore, inbreeding depression has negative effects both at the individual^30^ and population level^31^. Severe deleterious mutations were likely reduced through purging of highly deleterious mutations, while less severe mutations accumulated^5^. Previous genome-wide analyses of Alpine ibex showed that successful hybridization with domestic goats (*Capra hircus*) likely occurred during the severe historic bottleneck, when ibex numbers were at their lowest^32,33^. Introgression was particularly evident at immune-relevant genes, where genetic variability in Alpine ibex is very low. Such introgressed variants may have been under positive selection due to heterozygote advantage at immune loci^32^. However, hybridization is also expected to have detrimental consequences for Alpine ibex, with potential loss of adaptations to the alpine environment and the introduction of deleterious variants. Of particular concern are recently discovered hybrid swarms, populations of interbreeding hybrids between Alpine ibex and domestic goats in Northern Italy^27,34^. Such populations of hybrids have survived beyond the initial hybrid (F1) generation, and there is continued backcrossing between hybrid individuals and parental species^27^. This extensive hybridization raises concerns of maladaptation and loss of ibex identity.

**Figure 1:**
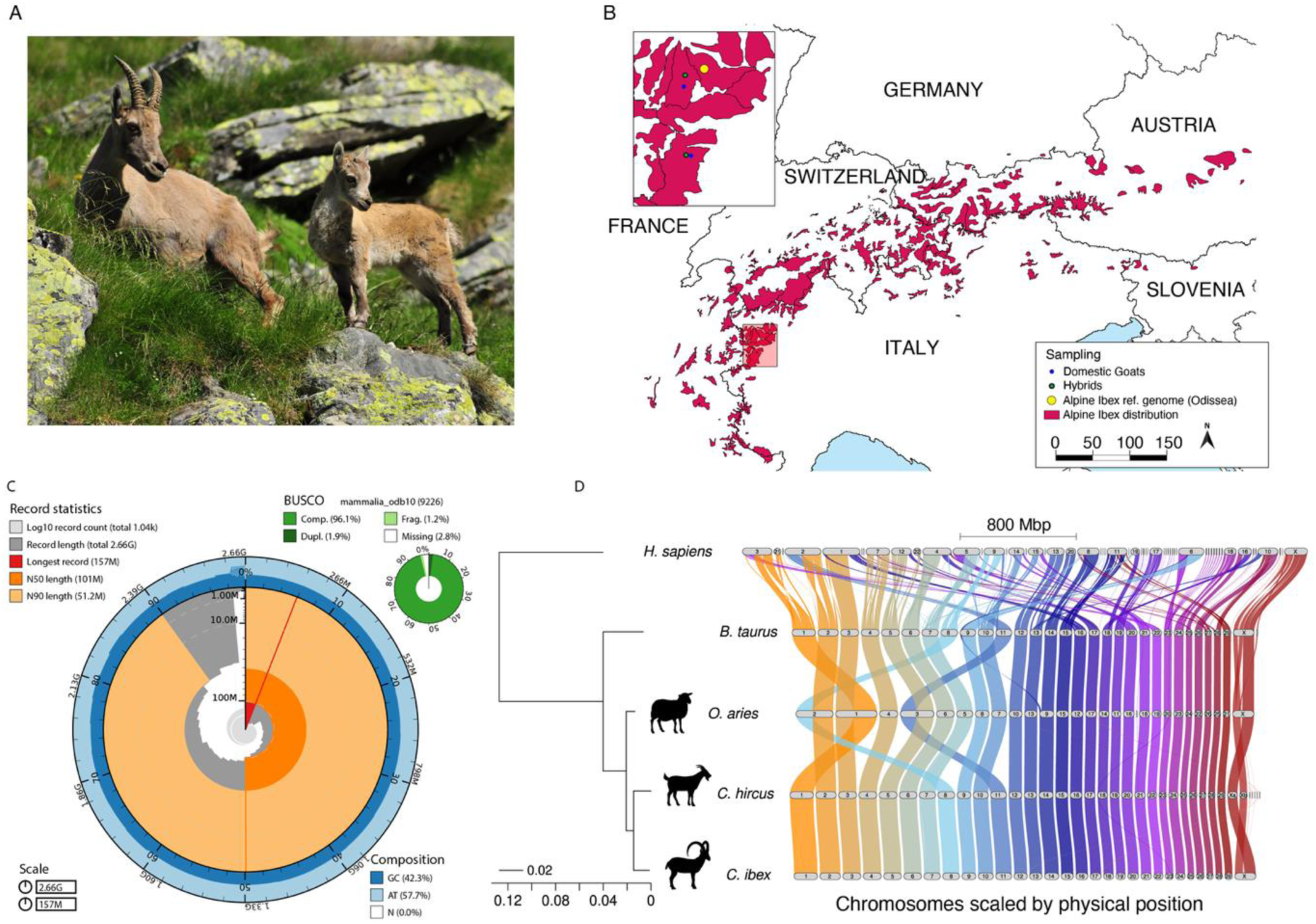
Sampling and genome description. A) Picture of a female Alpine ibex with her offspring (photo by Dario de Siena, Archivio PNGP). B) Map of species distribution and zoom in on the sampling location of Odissea, the female used for the genome assembly (shown in yellow), the hybrids (green), and domestic goats (blue). C) Snail plot visualizing assembly statistics: scaffold statistics, sequence composition proportions, and BUSCO completeness scores. D) Riparian plot showing the results of the gene synteny analysis conducted on the cattle, sheep, domestic goat, and Alpine ibex genomes using GENESPACE. The human genome was used as an outgroup. See also Figure S1 and Data S1.

Here, we generate the first reference-quality genome for Alpine ibex to quantify the hybridization load generated by high rates of hybridization in two hybrid swarms originating from interbreeding with domestic goats. We assess the consequences of introgression into Alpine ibex both at the population genomic level and through individual-based simulations to disentangle effects on pre-existing mutation load and disruption of local adaptation.

## Results and discussion

### A high-quality chromosome-level assembly

We assembled a chromosome-level genome of a wild female Alpine ibex (Odissea) sampled in the Gran Paradiso National Park population (Lauson Valley, Cogne, Italy). Using 130 Gb of high-quality ONT data, we generated a 2.66 Gb draft genome with 1040 contigs (N50 58.7 Mb), polished with short read sequencing data. The contigs showed high completeness, with BUSCO and k-mer-based scores of 96.1% and 98.6%, respectively (Figure 1C). We generated Hi-C data to anchor contigs into 30 pseudo-chromosomes (Figure S1) with a resulting N50 of 101.2 Mb and matching the chromosome numbers of domestic goats ^35^ (Data S1, Figure 1D). The Alpine ibex assembly also includes the full mitochondrial genome (16.7 Kb, Figure S1). The nuclear genome is constituted of 43.4% repetitive elements (1.15 Gb), dominated by retroelements (38.22%), including Long Interspersed Nuclear Elements (LINEs: 32.56%) and Long Terminal Repeats (LTRs: 4.87%) (Data S1). We integrated gene model evidence using transcriptome-assisted predictions as well as annotations lifted over from the sheep (*Ovis aries*) genome, yielding a BUSCO completeness of 99.7% at the transcript level (Data S1). The annotated gene set consisted of 32,783 protein-coding genes and 61,420 transcripts (Data S1).

### Recent structural variants arising in Alpine ibex

We used gene order conservation analyses between Alpine ibex and three related domesticated mammals, including cattle (*Bos taurus*), sheep, and domestic goat, as well as the human (*Homo sapiens*) genome as an outgroup to detect chromosomal rearrangements (Figure 1D; Data S1). Chromosomal synteny is largely conserved between the domestic goat and the Alpine ibex. The sheep genome experienced three chromosome fusions, combining in each two distinct chromosomes found in cattle and goats. The Alpine ibex genome includes a fully assembled X chromosome compared to the domestic goat (contigs Xa and Xb; Figure 1D). In addition, the Alpine ibex chromosomes 12 and 17 also include sequences matching to unplaced contigs in domestic goat. Overall, domestic goat and Alpine ibex genomes share 2.4 Gb of syntenic regions (∼90.2%). Structural variant analyses with SyRI confirmed major rearrangements on chromosomes 7, 18, and 23 (Figure S1), mainly involving indels (38 Mb, 1.4% of the genome) and inversions (49 Mb, 1.8%), with fewer duplications and translocations (Figure S1). Highly divergent regions (HDR) covered a total of 43 Mb (1.6%; Figure S1). Nearly half (45%) of all insertions, deletions, and HDRs between domestic goat and Alpine ibex overlapped with gene bodies, and the proportion was even higher for inversions (52%), duplications (54%), and translocations (78%).

### MHC conservation and high heterozygosity in recent hybrids

The major histocompatibility complex (MHC) is known for high within-species polymorphisms, and the locus is prone to duplications ^36^. Consequently, assembling the MHC regions can be challenging [see *e.g.,* challenges for human MHC assembly ^37–39^]. Here, we report a contiguous assembly of the Alpine ibex MHC. High-level synteny is well-conserved among cattle, sheep, domestic goat, and Alpine ibex in both their main MHC regions (class I and II, Figure 2A). We detected a ∼110 kb translocation between Alpine ibex and the domestic goat near the MHC class II cluster that does not affect MHC coding sequences (Figure 2B). An additional translocation of ∼1.92 Mb occurred further away from the MHC class II on chromosome 23 (Figure 2B). In contrast to the human MHC, both the domestic goat and Alpine ibex MHC class II clusters encode DR- and DQ-genes but no DP-genes ^40,41^. The DR- and DQ-genes are generally exceptionally polymorphic. Over 100 DRB1 alleles for sheep and dozens for goats have been deposited in GenBank, reflecting their crucial role in immune diversity. Alpine ibex populations have previously been shown to carry introgressed alleles from the domestic goat at the DRB gene ^33^ and other immune-related genes ^32^.

**Figure 2:**
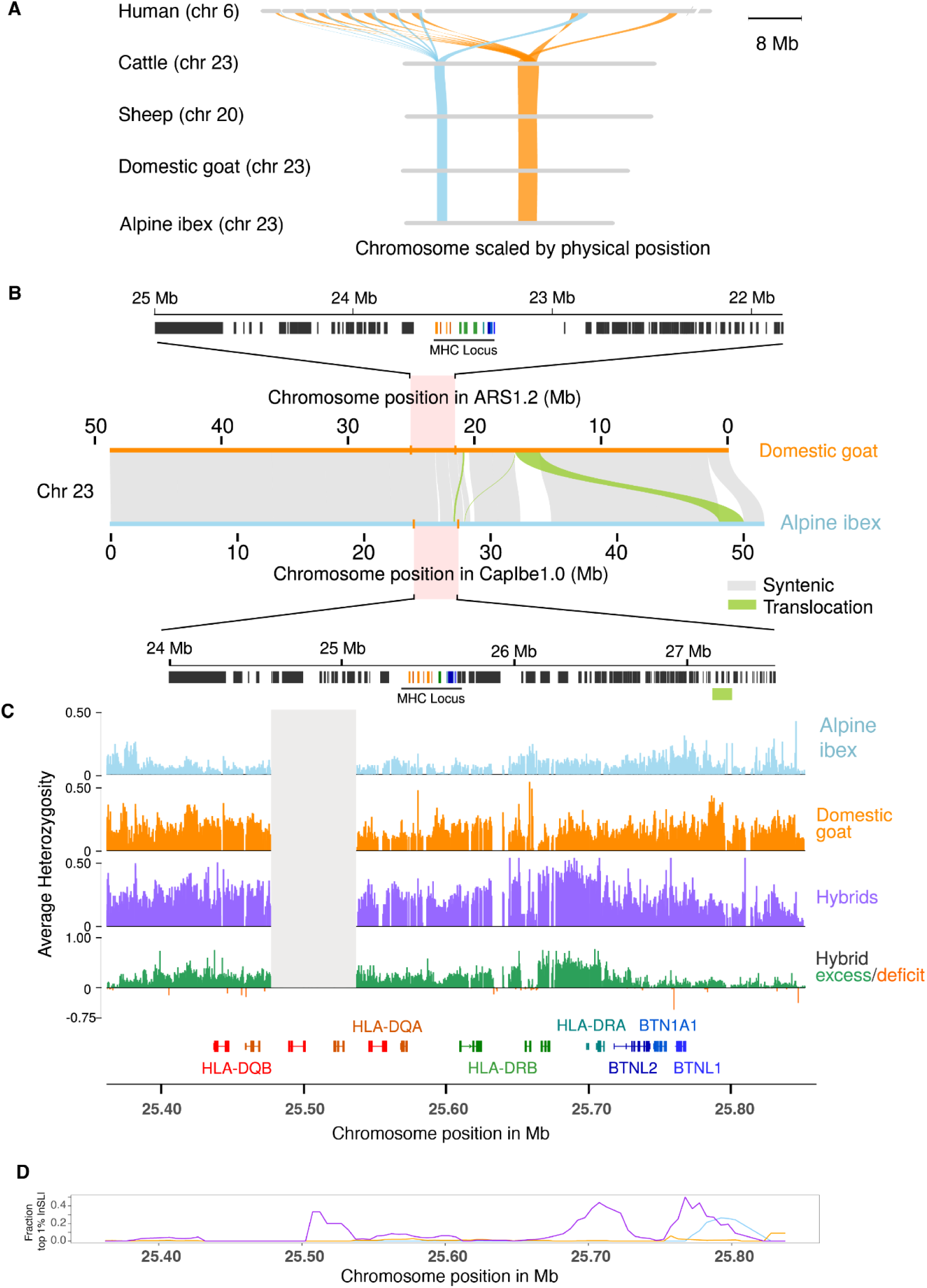
Conserved synteny at the MHC and high diversity in hybrids. A) Close-up of the synteny analysis, conducted with GENESPACE, highlighting two MHC regions within Alpine ibex chromosome 23 (light blue and orange, respectively). B) Structural variation along chromosome 23 based on SyRI, including annotations of domestic goat (top) and Alpine ibex (bottom) in the range encompassing the major MHC II region. C) Average individual heterozygosity is shown as observed heterozygosity in Alpine ibex (blue), domestic goat (orange), and hybrids (purple). Heterozygosity excess and deficit in hybrids are shown with green (positive values) and orange (negative values), respectively. The grey box highlights a region corresponding to 25.47-25.55 Mb (encoding *DQA* and *DQB*), which showed a local lack of coverage in domestic goats while being evenly covered in Alpine ibex (except for Alpine ibex individuals with previously reported historic introgression in the MHC region, see Grossen et al. ^33^). D) Local enrichment of unusually long haplotypes (extreme |nSL| values) across the MHC region on chromosome 23 was shown in the fraction of SNPs exceeding the genome-wide 99th percentile of normalized |nSL| in sliding windows of 25 Kb (5 Kb steps) across positions 25.35–25.85 Mb. Curves represent Alpine ibex (light blue), domestic goat (orange), and ibex– goat hybrids (purple). See also Figure S2 and Data S2.

To assess whether hybridization load may pose a risk to Alpine ibex, we investigated two recently discovered hybrid swarms in Northern Italy. One discovery followed a reportedly abandoned goat herd in the late 1990s in the same region ^34^. We generated whole-genome sequencing data of eight recent hybrid individuals (including backcrosses) ^27^ and four domestic goats representing the local breed, and collected in the same areas as the hybrids (Data S1). We analysed the new sequencing data together with publicly available data representing 29 non-hybrid Alpine ibex as well as 18 domestic goats. Average individual heterozygosity was consistently low in non-hybrid Alpine ibex individuals compared to hybrids and domestic goats, including particularly striking contrasts in the DR- and DQ-gene regions (25.40-25.75 Mb; Figure 2C). Increased diversity was also reflected in patterns of copy number variation within the MHC region, where hybrids (Figure S2) exhibit a mix of copy-number patterns seen either in domestic goat (Figure S2) or Alpine ibex (Figure S2). We also analyzed haplotypes at the DRA and DRB loci. Hybrids each carried a native Alpine ibex and a goat DR-allele (Figure S2). The increase of diversity was particularly striking at the DRB gene, where each of the eight hybrids gained a distinct domestic goat allele (Figure S2) creating opportunities for MHC introgression beyond the previously detected introgressed allele ^33^.

Diversity at the MHC is associated with disease resistance in a number of species ^42–48^, including Alpine ibex ^49^. The high diversity at the MHC in hybrids may therefore confer a selective advantage under disease pressure. We found support for this selection pressure through signals in selective sweep scans (Figure 2D, Figure S2). Hence, hybridization and MHC introgression may be favored in the wild, posing a risk of disrupting local adaptation in wild Alpine ibex.

### Emerging recombinant haplotypes in an Alpine ibex hybrid swarm

The recent observation of hybrid swarms in Northern Italy renewed concerns about Alpine ibex conservation efforts ^27^. The high collinearity of Alpine ibex and domestic goat genomes should facilitate recombination between haplotypes. At the phenotypic level, recent hybrids carry well-recognizable domestic goat features such as horn shape and fur coloration ^27^ (Figure 3A, B). We analyzed the whole-genome sequencing data to recapitulate introgression patterns in two hybrid swarms. We furthermore sequenced four domestic goat individuals from the same two regions (Figure 3, Data S1). Non-hybrid individuals of each species showed clear separation based on a principal component analysis (PCA, *n* = 29 Alpine ibex, *n* = 22 domestic goats; Figure 3C) performed on 86,650 biallelic SNPs. All eight hybrid swarm members showed clear evidence for recent admixture (Figure 3C). Next, we performed chromosome painting using genome-wide polymorphism of the hybrids to identify parental contributions. Individual 12 matched expectations for a F1 hybrid with uninterrupted goat and Alpine ibex haplotypes over all chromosomes (Figure 3D). Hybrids that underwent backcrosses revealed evidence for recombination events as interrupted haplotypes (Figure 3D). Genome-wide ancestry matched well with recombination patterns and parental haplotype contributions. Individual 5 showed both the highest degree of ancestry and the largest haplotypes of domestic goat. In contrast, individual 8 carried the longest Alpine ibex haplotype blocks consistent with its strong Alpine ibex ancestry (Figure 3D). The highly diverse set of recombinants in the hybrid swarms raises significant concerns for further introgression in the two respective regions and in other populations with a high incidence of contacts between domestic goat and Alpine ibex.

**Figure 3:**
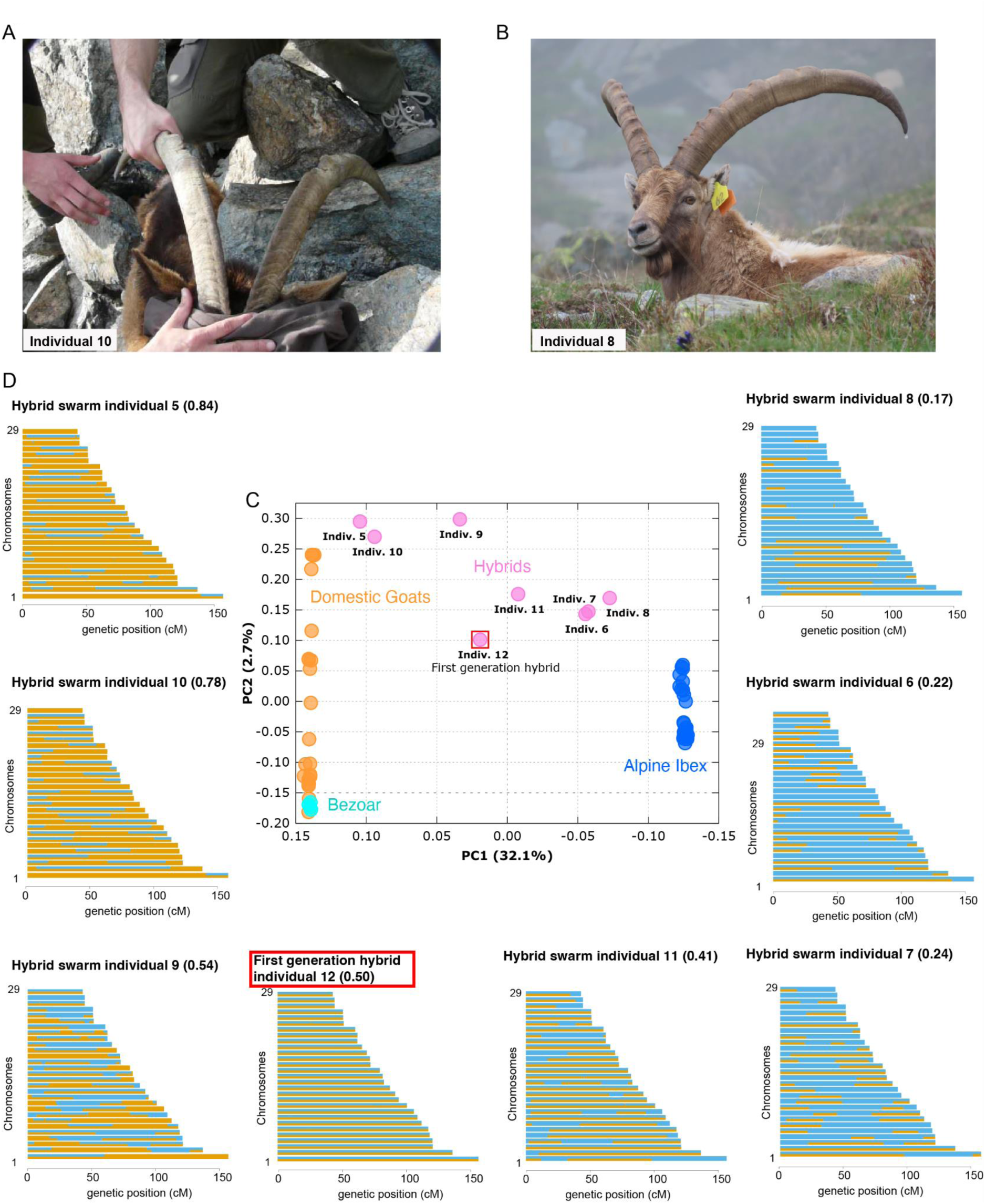
Genome-wide patterns of hybridization. A) Picture of hybrid individual 10. B) Picture of hybrid individual 8. C) Principal Component Analysis of biallelic autosomal SNPs showing the clustering of species and the distribution of hybrids along the conjunction between Alpine ibex and domestic goat/bezoar. D) Chromosome paintings drawn with Mosaic, ordered according to the proportion of domestic goat ancestry (decreasing proportion of goat genome shown in orange from left to right). Also see Data S1.

### Hybridization load in two Alpine ibex hybrid swarms

We assessed the extent of hybridization load, which could be introduced into Alpine ibex through a hybrid swarm. The cross-species polymorphism included 59,544,737 SNPs of which 0.009%, 0.311%, and 0.499% were classified as having putatively high, moderate, and low impacts on encoded proteins, respectively (see also Data S1). High-impact variants, such as those causing premature stop codons or frameshifts, are predicted to severely disrupt protein function (*e.g*., loss-of-function, LOF variants). Moderate and low impact variants cover, respectively, missense mutations that may alter protein structure without severe impairment and synonymous changes with minimal expected effects on protein function. To assess confidence in the predicted impact of mutations, we combined gene annotation with RNA-seq evidence of one Alpine ibex across 10 organs. We identified genes with annotated CDS and deemed them expressed and, hence, more likely functional if they had CPM ≥ 0.1 in ≥ 1 organ. With these filters, 69% of putative LOF variants fell into expressed CDS genes and were the only LOF mutations (N=88) retained for the genetic load analyses.

We investigated deleterious mutations derived from either domestic goat or Alpine ibex as a proxy for genetic load (Figure 4A). First, we analyzed candidate mutations underpinning heterosis in hybrids. We defined these mutations as fixed derived LOF variants in Alpine ibex (fixed for the ancestral allele in all other species). Only two LOF variants were identified as candidates for heterosis (Data S1). Severely disruptive variants such as LOF are generally recessive; hence, these should be masked in the heterozygous state, conferring increased fitness (heterosis) in hybrids compared to non-hybrid Alpine ibex. We found that the analyzed hybrids each carried between zero and both candidates for heterosis in the heterozygous state (in four backcrossed individuals and a F1 hybrid, respectively, Figure 4B). This suggests that the positive effects of masking LOF variants in hybrids may quickly get lost through backcrossing. The observed signs of hybrid vigour in Alpine ibex-domestic goat hybrids ^27^ may reflect a combination of such masking effects alongside other genetic (*e.g.*, overdominance) or epigenetic factors ^50^.

**Figure 4:**
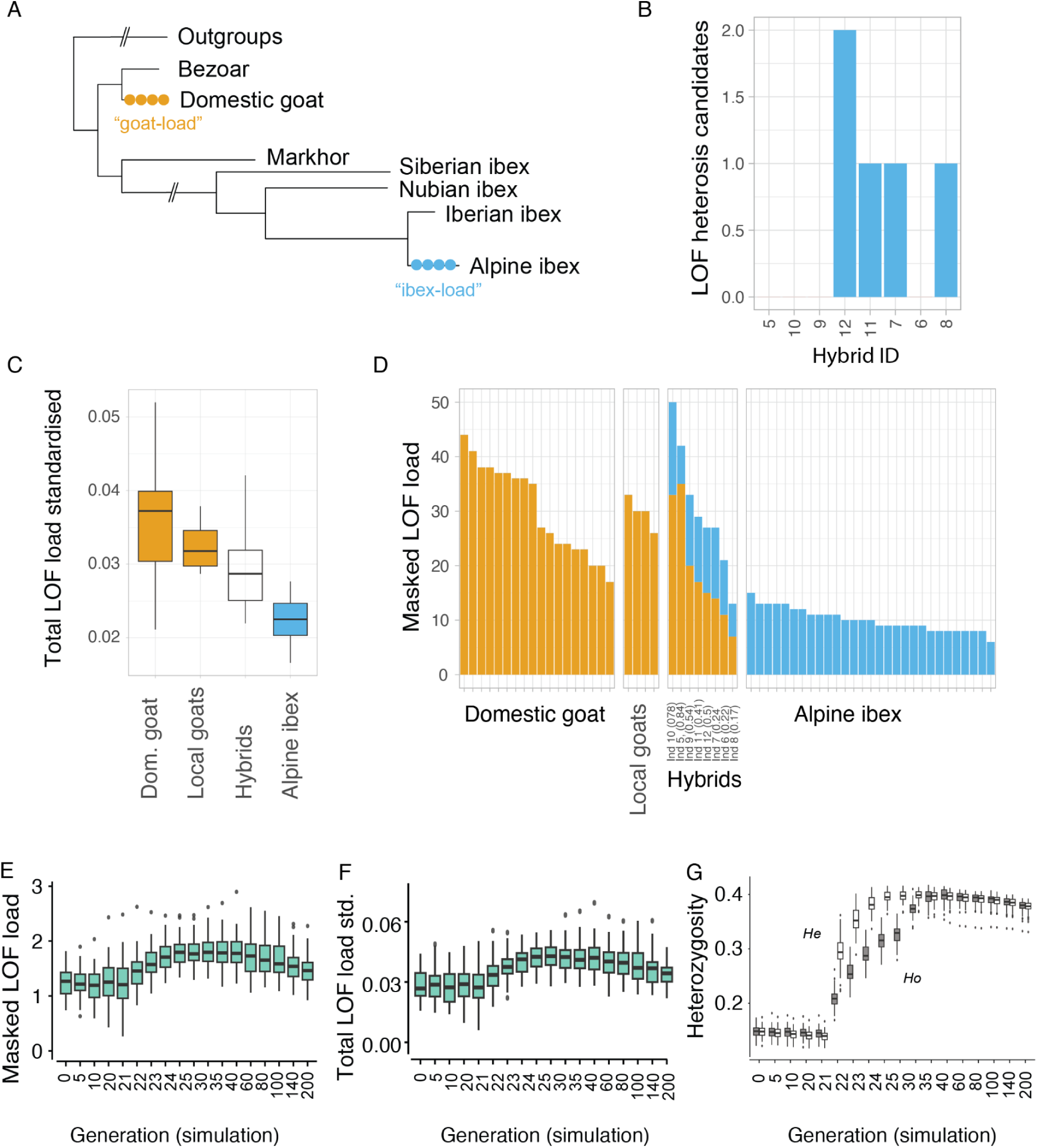
Genetic load in hybrids. A) Phylogenetic tree showing species relationships and the definition of “goat-load” and “ibex-load” defined as newly derived mutations private to the respective species. B) Absolute counts of loss-of-function alleles found in the heterozygous state for each sequenced hybrid individual (ordered according to proportion of goat ancestry). C) Standardized loss-of-function load (total counts of loss-of-function alleles standardized by the total counts of modifier alleles). Box plots per species. Local goats refer to domestic goat breeds kept in the region where hybrid swarms were detected. D) Masked loss-of-function load (absolute heterozygote counts) per individual. Local goats, as in C. E-G) Simulation results for 1 % dispersal pulse scenario: E) Masked loss-of-function load (absolute heterozygote counts), F) Standardized loss-of-function load (total counts of loss-of-function alleles standardized by neutral variation), and G) Genome-wide observed (H_o_) and expected (H_e_) heterozygosity (averaged over loci) across simulated ibex generations. Boxplots per generation show replicate (N=50) variation. Differences between observed and expected heterozygosity (*i.e.*, deviations from Hardy–Weinberg equilibrium) are expected due to immigration. See also Figure S3, Data S3.

We also investigated the extent to which hybridization load could affect Alpine ibex. We found that the standardized LOF load, which is normalized by neutral variation for cross-species comparisons, was higher in domestic goat than in Alpine ibex (Figure 4C, p < 10^−6^). Estimates of standardized LOF load were also higher in the closely related Iberian and Siberian ibex (*C. pyrenaica*, *C. sibirica*) compared to Alpine ibex (Figure S3); however, domestic goats are the most significant threat to Alpine ibex hybridization swamping given the overlap in distribution range. Consistent with these expectations, members of the Alpine ibex hybrid swarms carried a total of 88 newly acquired LOF variants within expressed CDS (Data S1), more than doubling the total number of LOF variants in Alpine ibex (n = 58 LOF variants detected among 29 non-hybrid Alpine ibex). This corresponds to an average of 30 masked LOF variants per hybrid individual compared to 10 in non-hybrid Alpine ibex. Hence hybrids carried on average approximately three times higher masked LOF load ^51^ (Figure 4D). Among the hybrids, those backcrossed with Alpine ibex saw their LOF load decreased with each generation as expected (Figure 4D). Yet standardized LOF load did not decrease over the first few generations of backcrossing into Alpine ibex (Figure S3). Hence, within the limited sample size, there is no indication that selection acted to reduce hybridization load. Despite the lack of direct fitness estimates for hybrids and backcrosses, the persistence and expansion of the hybrid swarms reported in Alpine ibex ^34^ suggest no major counter-selection. Backcrosses up to the 4th-5th generation ^27^ as well as pregnant female hybrids were observed repeatedly. Backcrossed males (*e.g.*, individuals 6, 7 and 11) tended to be large and with significant secondary sexual traits (horns and testicle size, Data S1). Hence, circumstantial evidence is consistent with no or only minimally reduced hybrid fitness. Consequently, further backcrossings (and recombination) will likely expose Alpine ibex to the risk of increasing their genetic load through leakage from the hybrid swarm.

To explore the consequences of different hybridisation scenarios (*i.e.*, different rates and durations of gene flow) between a genetically depleted species (*i.e.,* Alpine ibex) and a species with higher genetic diversity (*i.e.,* domestic goat), we performed individual-based genomic simulations (Figure 4E-G, Data S3). We assessed the consistency of our empirical results with predicted dynamics of hybridization load and purging. Masked LOF load (here defined as the sum of heterozygous LOF sites as a proxy) increased from an average of 1.2 LOF mutations per individual to an average of 1.8. Furthermore, standardized total LOF load increased by a factor of about 1.5. A higher rate of gene flow from domestic goat to Alpine ibex (increase from 1% to 2.5%) led to a faster increase in LOF load but only slightly higher overall levels (average of ∼2 LOF mutations, Data S3). Approximately 35 generations after the pulse of gene flow ended, masked LOF load decreased again and declined toward pre-hybridization levels over the subsequent 140 generations (or 175 generations after gene flow subsided) (Figure 4E). Such a reduction in LOF load was not observed under ongoing gene flow, which would lead to genetic swamping of the species (see Data S3). Interestingly, we observed a slight decrease of LOF load in ibex if no gene flow occurred (Data S3). While neutral genetic diversity (a proxy of adaptive genetic diversity^52^) increased substantially under gene flow (Figure 4G), it slowly decreased over time due to drift in the absence of gene flow (Data S3). Hence, our simulations support a model, in which hybridization can introduce LOF load into species with low diversity.

Importantly, we noted that mildly deleterious mutations accumulated in the simulated ibex population, while these were very efficiently removed by selection in the domestic goat. This explains the slight fitness benefit of ibex following gene flow from the domestic goat (Data S3). Accumulated mildly deleterious mutations are replaced in part by introgression. This slight fitness benefit is also consistent with a previous empirical study of Alpine ibex genomes, which suggested that preventing accumulation of mildly deleterious mutations may be overall more beneficial than purging few highly deleterious mutations ^5^. Beyond considerations of LOF, Alpine ibex may also be threatened by the disruption of local adaptation to the alpine environment by the introduction of maladaptive goat alleles. Simulations including a locally adapted trait substantiated this scenario and indicated a significant reduction in Alpine ibex fitness following hybridization (Data S3). Furthermore, balancing selection likely acting on MHC polymorphism could favor introgression by conferring immunogenetic benefits ^32,33^. In conjunction, our results show that fitness consequences for a vulnerable species exposed to a hybrid swarm are multifaceted and warrant further investigation.

## Conclusions

The detrimental effects of deleterious mutations are well-recognized as a risk factor in endangered species ^2,3^. While a large number of recent studies investigated the evolution of genetic load as a consequence of small population size ^4–6,53–57^, much less consideration was given to its source. Here, we investigated the role of hybridization as a source of genetic load. Hybridization produces genetic variation and can foster adaptation ^11,12,58^. Indeed, introgression from domestic goat into Alpine ibex has previously been suggested to be adaptive, in particular at immune-relevant genes ^32,33^. Yet high levels of hybridization raise concerns regarding potential detrimental consequences ^27^. A number of studies on fish ^14,15^, crops ^16–18^, and trees ^19^ show that although hybridization can introduce beneficial alleles, linked deleterious variants can reduce fitness. Furthermore, hybridization poses the risk of disrupting local adaptation through the introgression of non-native alleles as shown in our simulations. This risk is likely particularly acute if the donor species shares the recipient species’ habitat only because of domestication or other human activity ^20–23^. Such swamping of the native gene pool is particularly evident in Scottish wildcat, where hybridization has affected immunity-related functions ^24^.

Here, we show that high rates of hybridization produced by a hybrid swarm can introduce severely deleterious mutations from a closely related domestic species. The initial masking of deleterious mutations in hybrids is likely to reduce efficiency of selection against them in early generations, consistent with our observations. Access to a larger number of hybrids would enable genomic-tract based analyses and direct assessments of purging. High genome-wide heterozygosity in hybrids may further confer early-generation hybrids an advantage over non-hybrid individuals, compounding the risk of further introgression and a potential loss of local adaptation. In a genetic rescue scenario, the beneficial effects of increased diversity are expected to overcome possible detrimental effects of introduced LOF load as indicated by our simulations ^59–61^. However, anthropogenic hybridization and the risk of introducing hybridization load, as well as maladapted alleles can be seen as a form of genomic erosion ^62,63^. Given the increased rate of human-induced hybridization ^64^, comprehensively tracking fitness effects of hybridization and assessing long-term consequences for the gene pool of the vulnerable species should be a central concern of species conservation actions. Empirical data on hybrid fitness will be particularly valuable for validating these risks and informing management decisions.

## Data availability

Raw sequencing experiment data were submitted to the SRA archive under the project identifier PRJNA1104824 (https://www.ncbi.nlm.nih.gov/datasets/genome/GCA_054642885.1). The final *C. ibex* genome assembly and high-confidence coding gene annotations are available under the NCBI accession number GCA_054642885.1.

The *CapIbe1.0* genome fasta file, full transcriptome models, gene functional annotation, and GERP scores relative to the novel genome can be downloaded from Zenodo (https://doi.org/10.5281/zenodo.15632776; https://doi.org/10.5281/zenodo.15632854; https://doi.org/10.5281/zenodo.15357984).

## Code availability

Codes used for variant classification, load quantification, and individual-based simulations are available from GitHub: https://github.com/fgcilingir/ibex_hybrid_load

## Acknowledgments

For domestic goat sampling, we thank Stefania Zanet, Alice Naudin, and Emilio Gugliermetti. For hybrid sampling, we thank the Surveillance Service of Gran Paradiso National Park, the Forestry Service of Regione Valle d’Aosta, Claudio Scaini, and the personnel of Città Metropolitana di Torino and of CATO4, Noel Zehnder and Luca Rossi. We thank Sara Gottardo, Francesca Groppo, and the sequencing facility of CMP^3^VdA in Aosta for technical assistance with Illumina whole-genome sequencing. We thank Sergio Decherchi, Alessandro Parodi, and Mattia Pini for their support for high-performance computing and data storage, and Dario de Siena and Raffaele Turvani for the beautiful ibex pictures. We thank Fred Guillaume for very helpful advice regarding the simulations and Iris Biebach for guidance on Alpine ibex age structure.

## Funding information

F. Gözde Çilingir was funded by the Swiss National Science Foundation (IZCOZ0_198147 to A.B. and C.G. and 31003A_182343 to C.G.).

This work was supported by the 5000genomi@VdA Project co-founded by “Fondo Europeo di Sviluppo Regionale (FESR CUPB68H19005520007) to SG and AC; FL, ST, ACa, MV and LP were supported by “Fondo Europeo di Sviluppo Regionale (FESR CUPB68H19005520007); DC and FF were recipient of a fellowship from Programma Investimenti per la crescita e l’occupazione 2014/20” (European Social Fund, ESF CUP B65F19001200009) and continued their activity with a fellowship from Programma Regionale Fondo Sociale Europeo Plus 2021/2027 (“European Social Fund”, ESF CUPJ51B24000170002).

This work is part of the “Technologies for Sustainability” Flagship program of IIT (L.P.).

## Author contributions

AB conceived the study and managed DNA sample collection. AB, AP, GC, DW, JHM, and BB performed and coordinated the sampling. FGÇ, FL, DC, LP, and CG designed experiments and analyses. EH, YCCDS, and DV generated sequencing data under the supervision of FGÇ, LP, MV, and ST. FGÇ, FL, FF, LP, and CG performed analyses with input from DC. FGÇ performed the simulations with input from CG and DC. FGÇ, FL, AB, DC, LP, and CG wrote the manuscript with contributions from all authors. JMC, ACa, SG, AB, and CG provided funding and resources for the study.

## Material and methods

### Biological material

Fresh samples of different organs (spleen, muscles, liver, kidney) were collected from a wild female Alpine ibex from the population of Gran Paradiso in the North-Western Italian Alps (Figure 1B). This individual was sampled in an area where no feral goats have been observed in the recent past and it had a typical Alpine ibex phenotype without any suspicious phenotypic trait that could indicate recent hybridization. Genotyping at 63 diagnostic markers as described in ^27,65^ confirmed that this individual was not a recent hybrid. Immediately after the animal died of natural causes in Lauson Valley, Cogne (AO, Italy) on 18 June 2021, it was brought to the laboratory of Gran Paradiso National Park and was immediately dissected.

For the hybrid studies, blood or tissue samples were collected from eight hybrid individuals (with different levels of expected hybridization) between Alpine ibex and domestic goats, and four domestic goats representing local breeds (Data S1). The samples were collected within the framework of an extensive sampling campaign carried out in two hotspots of suspected hybrid presence^27,34^: Gran Paradiso National Park (Valsavarenche, Aosta, Italy) and Lanzo Valley (Torino, Italy). Suspected hybrids were originally identified based on phenotypic characteristics and either captured with chemical immobilization or culled, and their hybrid status (F1 and backcrosses) was subsequently confirmed based on 63 diagnostic SNPs ^27^. Domestic goat samples were collected in the same areas where the suspected hybrids were observed, during routine sanitary monitoring of the domestic herds.

### DNA extraction for genome assembly

The samples for the genome assembly were collected within four hours of the animal’s death, placed in cryo-stable plastic tubes (three aliquots from each organ), and immediately frozen at −80°C. High molecular weight (HMW) DNA extraction was performed to allow long-read sequencing for d*e novo* assembly of the reference genome of Alpine ibex. Four separate extractions from spleen (N=2) and liver (N=2) tissue were carried out with the MagAttract HMW DNA Kit (QIAGEN). An additional fifth extraction with spleen tissue was performed using the Monarch® HMW DNA Kit (New England BioLabs) for Ultra-Long library preparation.

### DNA extraction for hybrid studies

Blood samples were collected in EDTA vacutainer tubes. Tissue samples were stored in absolute ethanol at room temperature until DNA extraction, which was performed using QIAmp DNA mini and Biosprint 96 systems (QIAGEN), applying the appropriate protocol for each type of sample.

### Short-read library preparation and sequencing

Whole genome sequencing of gDNA (representing the reference individual, eight hybrids, and four domestic goats) was performed on an Illumina NovaSeq 6000 platform (Illumina Inc., CA, USA) to generate paired-end 151 bp reads. Individually barcoded hybrids were pooled during library preparation and sequenced together, whereas the library preparation and sequencing of the reference individual’s gDNA were performed separately. For all samples, 350 ng of gDNA were used for library preparation with Illumina DNA PCR-Free Prep, Tagmentation library preparation kit (Cat number 20041795), and IDT® for Illumina® DNA/RNA Unique Dual Indexes Set A or B, Tagmentation (96 Indexes, Cat number 20027214). Libraries were pooled by volume to reach a 90X coverage for the Alpine Ibex reference genome and a 25X coverage for hybrid samples. The Illumina bcl2fastq2 software was used to convert the original raw data format (bcl) to fastq.gz read files. Initial quality checks (including coverage and base qualities) were performed using FastQC ^66^.

### Long read library preparation and sequencing

For long-read sequencing, PromethION 24 instrument (PRO-PRC024) was used following the instructions from Oxford Nanopore Technologies (Oxford, UK). The R9.4.1 chemistry flowcells (FLO-PRO002) was used with multiple library preparation kits, including Ligation Sequencing Kit (SQK-LSK110), Rapid Sequencing Kit (SQK-RAD004) and Ultra-Long Sequencing Kit (SQK-ULK001). Reads were basecalled using Guppy version 6.1.1 in Super-High Accuracy mode (using the configuration dna_r9.4.1_450bps_sup_prom.cfg). Only reads with a Q-score greater than 10 were retained for subsequent analysis. For the final yield of each run see Data S1.

### Hi-C sequencing

The Hi-C sequencing library was prepared using 500 mg of liver tissue. Initially, the liver sample was treated with 1% formaldehyde for 15 minutes at room temperature for fixation. Next, solid glycine powder was added to achieve a final concentration of 125 mM, and the mixture was incubated for 15 minutes at room temperature with intermittent mixing. After centrifugation, the resulting pellet was resuspended in PBS + 1% Triton-X solution, followed by a 15-minute incubation at room temperature. The tissue was then collected by centrifugation of the mixture, transferred to a liquid nitrogen-cooled mortar, and ground to a fine powder. Finally, the cross-linked sample was shipped to Phase Genomics (Seattle, Washington, USA), while kept frozen on dry ice for sequencing. The Hi-C library construction was performed using the Phase Genomics Proximo Animal kit version 4.0. In summary, the DNA sample was digested with a fragmentation enzyme mix (including enzymes DpnII, DdeI, HinfI, and MseI), the 5’-overhangs were filled with a biotinylated nucleotide, the resulting fragments with blunt ends were ligated, and then sheared. The biotinylated ligation junctions were captured using streptavidin beads. Finally, the resulting DNA fragments were sequenced using an Illumina NovaSeq 6000 instrument (Illumina Inc., CA, USA) with a 150 bp paired-end run.

### Genome assembly and polishing

The ONT sequencing data were assembled into the draft genome using the software Flye v2.9 ^67^ with parameter “*`--nano-hq`*”. Then, the short-read data were aligned to this initial draft assembly using the GPU-accelerated fastq2bam tool of the NVIDIA Clara Parabricks v3.8 suite ^68^ with default settings. The tool performs the following steps: alignment with BWA-mem (v0.7.15) ^69^; sorting the coordinates with SortSam (GATK v4.1.0 ^70,71^); and finally, applying MarkDuplicates (GATK v4.1.0). The resulting aligned data were then used to polish the assembly with Pilon v1.23 ^72^.

### Contamination Scan

In order to identify and remove potential contaminant sequences in the assembled genome, the tool Foreign Contamination Screen (FCS) developed by NCBI ^73^ was used. The genome was scanned initially with FCS-adaptor to identify short foreign sequences coming from laboratory processes and subsequently with FCS-GX ^73^ to detect contamination from foreign organisms coming from environmental sources. Two small isolated contigs (3.9 and 47.8 Kb, respectively) representing putative reverse-transcribed viral RNA were removed.

### Genome scaffolding using Hi-C sequencing data

Hi-C sequencing reads were quality filtered with Trimmomatic v0.39 ^74^ with parameters “`SLIDINGWINDOW:30:10 LEADING:19 TRAILING:19 AVGQUAL:20 MINLEN:30`”. Quality-filtered reads were mapped on the draft Alpine ibex genome by following the Arima Genomics mapping pipeline (https://github.com/ArimaGenomics/mapping_pipeline). Briefly, reads were mapped in single-end mode using BWA-MEM v0.7.17 ^69^, chimeric joins across ligation junctions were filtered, and lastly, PCR duplicates were removed with Picard v2.6.0 ^75^. In total, 99.5% of the Hi-C reads aligned to the draft genome assembly and ca. 88% of the reads were found to have valid Hi-C contacts. The alignment files were then incorporated into the genome scaffolding using YaHS v1.1 ^76^ with default settings. The scaffolded contigs were examined visually within JBAT v2.10.01 ^77^ (Figure S1), and no manual refinement was found necessary.

### Assembly quality evaluation

Main genome contiguity statistics were estimated with QUAST v5.0.2 ^78^. The k-mer completeness analysis and the reference-free QV estimation were performed with merqury v1.3 ^79^ using the k-mer multiplicity of Illumina WGS reads utilized for the polishing of the Alpine ibex draft genome (see the section for Library Preparation and Sequencing of Short Reads) at k=21. The completeness of the assembly was also assessed based on evolutionarily informed expectations of gene content from near-universal single-copy orthologs via BUSCO v5.4.4 ^80^ with default parameters, using the mammalian dataset (mammalia_odb10) in the genome mode. Genome assembly statistics were visualized with a snail plot using BlobToolKit v4.3.2 ^81^ (Figure 1C).

### Repeat annotation and masking

A species-specific *de novo* repeat library for the Alpine ibex genome was generated using RepeatModeler v2.0.3 ^82^. For this, a species-specific repeat database was created with the BuildDatabase function of RepeatModeler using NCBI as the search engine. Then, *de novo* repeat family identification and modeling was performed with RepeatModeler utilizing RECON v1.08 ^83^, RepeatScout v1.0.6 ^84^, Tandem Repeats Finder (TRF) v4.09.1 ^85^, and RMBlast v2.11.0 ^86^ on blast databases formatted with BuildDatabase. These consensus sequences were then used to softmask the genome with RepeatMasker v4.1.2 (https://www.repeatmasker.org/) using “`-xsmall`” flag. Additional masking by TRF ^85^ with maximum repeat period size of 500 was applied in order not to skip long tandem repeats since the default run of RepeatModeler does not identify tandem repeats with repeat pattern length of more than 10. Finally, merging of the repeat coordinates obtained from RepeatMasker outputs and additional masking with TRF was performed, and the softmasking of the genome with these merged coordinates was applied with bedtools v2.30.0 ^87^.

### Gene model prediction, annotation, and curation

The gene model prediction was performed by using transcriptome and protein-homology-based approaches. Evidence for the transcriptome-based prediction was based on the already available RNA-seq dataset generated from retina/uvea, skin, heart, lung, lymph, bladder, ovary, kidney, liver, and spleen tissues of a single, non-hybrid Alpine ibex individual (NCBI BioProject PRJNA517635 ^5^). The scaffolded Alpine ibex genome was indexed with STAR v2.7.2a ^88^, and the RNA-seq data were aligned to the indexed genome using the rna_fq2bam tool of NVIDIA Clara Parabricks v3.8 ^68^.

The evidence for the protein-homology-based prediction was produced by combining all vertebrate proteins in the OrthoDB v10 ^89^. This dataset was aligned against the softmasked, chromosome-level assembled reference genome via the ProtHint pipeline v2.6.0 ^90–94^.

RNA-seq and homology-based evidence were incorporated for the braker2 v2.1.6 pipeline separately by running the software with “`--etmode`” and “`--epmode`”, respectively ^91,95–102^ with pre-trained parameter sets for human (*Homo sapiens*), which is the evolutionarily closest taxon for Alpine ibex available within the software. Finally, all gene models derived from these runs were integrated into a high-confidence, non-redundant gene set using TSEBRA v1.1.1 ^103^ with the default configuration set.

Consequently, an annotation lift-over from the sheep (*Ovis aries*) reference genome (ARS-UI_Ramb_v2.0, NCBI Genome GCF_016772045.1 ^104,105^) was conducted with Liftoff v1.6.3 ^106^ with “`-copies`” and “`-polish`” options. Following this approach, extra copies of genes that were not annotated in the reference after the initial lift-over were searched, and the exons were realigned for polishing via restoring proper coding sequences in cases where the lift-over resulted in start/stop codon loss or introduced an in-frame stop codon. Finally, gene model prediction and lift-over results were merged with TSEBRA ^103^, using “`--keep_gtf`” option and the default configuration set.

In order to reduce the redundancy of the transcript set obtained, we employed an isoform pruning pipeline as previously described ^107^. Briefly, transcripts were grouped according to the genomic locus position using gffread v0.12.7 ^108^. The longest Open Reading Frame (ORF) from each transcript was predicted using TransDecoder v5.5.0, and any shorter putative ORFs were excluded from further consideration. ORF clustering was performed using CD-HIT v4.8.1 ^109^ with a 90% identity threshold to group transcripts into clusters based on sequence similarity. Among the transcripts clustered within each gene, the top two transcripts with the longest ORFs were retained for each CD-HIT cluster. Genes lacking predicted ORFs in any of their transcripts were identified and separated into a distinct file. This step was implemented to safeguard very short monoexonic genes or non-coding RNAs (ncRNAs) from being overlooked during subsequent analyses. Finally, gene models were annotated using eggNOG mapper v2.1.11 ^110^ in DIAMOND-search mode.

The completeness of the final annotation was assessed based on single-copy orthologs via BUSCO v5.4.4 ^80^ with default parameters and the mammalian dataset (mammalia_odb10) in the protein mode.

### Synteny analyses and structural rearrangement identification

We used GENESPACE v1.2.3 ^111^ with default parameters to investigate synteny between human (GRCh38.p14, NCBI RefSeq GCF_000001405.40), cattle (ARS-UCD1.3, NCBI RefSeq GCF_002263795.2), sheep (ARS-UI_Ramb_v2.0, NCBI RefSeq GCF_016772045.2), domestic goat (ARS1.2, NCBI RefSeq GCF_001704415.2), and the Alpine ibex genomes. The transcript isoform corresponding to the longest ORF for each gene in a genome annotation was extracted. Only primary chromosome scaffolds were taken into account. Riparian plots were generated, setting the domestic goat genome as a reference. Further, syntenic regions and genome-wide rearrangements were identified only between the closely related domestic goat (ARS1.2, NCBI RefSeq GCF_001704415.2) and Alpine ibex genomes. For this, all chromosomes from each genome except for the sex chromosomes were aligned to each other using minimap2 v2.26 ^112^ with default parameters (“`-ax asm5`”). The resulting alignments were processed with SyRI v1.6.3 ^113^ to identify syntenic regions and structural rearrangements. The syntenic regions and structural rearrangements were visualized with plotsr v1.1.1 ^114^.

### SNP genotyping

A dataset was compiled comprising the newly produced whole-genome re-sequencing data from eight hybrids and four domestic goats and available WGS data of Alpine ibex (N=29), domestic goat (N=18), bezoar (*Capra aegagrus,* N=4), Iberian ibex (*Capra pyrenaica,* N=4), Nubian ibex (*Capra nubiana,* N=2), Siberian ibex (*Capra sibirica,* N=2), and markhor (*Capra falconeri,* N=1). Additionally, four individuals representing outgroup species (chamois, *Rupicapra rupicapra*, cattle, *Bos taurus,* sheep, and the bighorn sheep, *Ovis canadensis*) were downloaded (Data S1). The downloaded WGS data were aligned to the newly generated chromosome-level assembly of Alpine ibex using the *germline* pipeline of the NVIDIA Clara Parabricks v3.8 ^68^, activating the flag “*`-gvcf`”* and without specifying a reference data set (no “*`--known-indels`”* option). The pipeline initially executed the fastq2bam tools (as described in the section “Genome assembly and polishing”). Then a GPU-accelerated version of HaplotypeCaller (GATK v4.1.0) for the variant calling was utilized. Later, a consolidation step was performed, where all the GVCFs generated in the previous step were merged into a cohort file with CombineGVCFs (GATK v4.1.0). To speed-up the execution, the genome was divided into intervals of similar size (each interval could contain one or multiple contigs). Each cohort GVCF was then used for the joint genotyping with GenotypeGVCFs (GATK v4.1.0). Finally, the compressed VCF obtained from each region was merged into a single file containing all SNPs and indels from all samples.

The VCF containing all variants underwent a hard-filtering process: first, quality flags were set for each variant call using the function VariantFiltation (GATK v4.1.0) with thresholds set to “`QD>=2.0, FS<=40.0, SOR<=4.0, MQ>=30.0, MQRankSum>=-4.0, MQRankSumH<=4.0, ReadPosRankSum>=-2.0, ReadPosRankSumH<=2.0`” and “`AN>=37`”. Subsequently, the variants that did not match these quality standards were filtered out using the SelectVariants function (GATK v4.1.0). We then extracted all SNPs polymorphic among the *Capra spp.* (removing all SNPs only polymorphic among the outgroup) which were autosomal, bi-allelic, with a minimal genotype quality of 20 and a minimal genotyping rate of 80%, using bcftools v1.2 ^115^. Average observed heterozygosity per-site was estimated for each species/group using the `-hardy`” option of VCFtools ^116^.

### Principal component analysis

The SNPs polymorphic within the *Capra spp.* were used to perform a Principal Component Analysis (PCA) with PLINK v2.0 ^117^. All variants with minor allele frequency below 0.05 and genotype missingness rate of over 10% were discarded (“`--maf 0.05 --geno 0.1`”). Then, the variants in linkage-disequilibrium were filtered out (“*`--indep-pairwise 1000 200 0.2`”*). The variants that passed these filters were used to perform the PCA. The first two principal components explained 34.8% of the data variation (respectively PC_1, 32.1% and PC_2, 2.7% Figure 3C).

### GERP scores for the C. ibex genome

Genomic Evolutionary Rate Profiling (GERP) scores of the *C. hircus* (ARS1) genome were lifted over to the *C. ibex* genome. Briefly, the two genomes were aligned using minimap2 ^112^ (with flags “`-cx asm5 --cs`”), then the output was converted into a chain file by paf2chain ^118^. GERP scores were lifted over after an intermediate bedgraph file conversion and saved back into bigWig format using the tools bigWigToBedGraph, liftOver, and bedGraphToBigWig UCSC Kent utilities ^119^.

### Chromosome painting

To analyze Alpine ibex and domestic goat haplotypes (and perform chromosome painting), the R-toolbox Mosaic v1.5.1 ^120^ was used. Mosaic searches for haplotypes that are shared between defined target individuals (here eight hybrids and four domestic goats) and individuals in several putative source panels (here domestic goat, bezoar -the wild ancestor of domestic goat, *C. aegagrus*-, Iberian ibex, Siberian ibex, and Nubian ibex, *C. nubiana*). Then the tool builds an admixture model, which is consistent with mixing between unseen groups giving rise to the observed data. Before running Mosaic, VCFtools ^116^ was used to thin the data down to a minimal inter-SNP distance of 1 Kb. Next, Phasing was performed using Beagle v22Jul22.46e ^121^, assuming a linear relationship between genetic and physical distance (1 Mb = 1 cM) because no recombination map is available for Alpine ibex. Genotype files per group were created using VCFtools ^116^ using the option —keep with the respective sample lists and SNP files using the option —plink before running Mosaic in R v4.3.2 ^122^. Chromosome painting visualization was performed using the function karyogram in Mosaic.

### Mutation load analysis

SnpEff v4.3t ^123^ was used to annotate variants’ putative functional changes. For this, the structural gene annotation generated was employed along with the bi-allelic SNP dataset containing polymorphic variants within *Capra* spp., (see section “SNP Genotyping”). The genome annotation was refined by filtering out transcripts containing in-frame stop codons using gffread v0.12.7 (option -V) ^108^. The longest transcript was retained in cases of overlapping transcripts for the same variant using the agat_sp_keep_longest_isoform.pl script within AGAT v1.4.1 ^124^. SnpEff was run using default settings to build a database with the new reference genome and the refined genome annotation. Then, variants were annotated across intergenic and genic regions retrieving putative effect categories of high (*e.g*., stop loss, start loss, frameshift variants), moderate (*e.g.*, missense variant, 3’ and 5’ UTR variant), low (*e.g.*, synonymous variant), and modifier (*e.g.*, intron variant, intragenic variant). Sites annotated as high (considering only the primary annotation) and containing a LOF variant were defined as putative LOF sites. As a first step in assessing robustness of LOF predictions, we analyzed the degree of gene conservation by searching for encoded PFAM domains. For this, the snpEff-annotated variant file was pre-processed using Unix command-line tools to extract transcript-level, protein-coding annotations and retain only the fields describing effect, impact, gene, and transcript IDs. The resulting table was imported into R ^122^, where each variant was annotated as PFAM+ or PFAM based on whether the encoded protein sequence contained PFAM domain hits. Variants were then grouped by SnpEff impact category and PFAM status to calculate the mean number of mutations per transcript. We postulated that LOF should occur also in conserved CDS (*i.e*., encoding PFAM domains). A bias towards PFAM-would have indicated a potential bias towards calling LOF in non-conserved regions. But we found no significant difference between proportions of LOF in CDS encoding PFAM domains (*i.e.* conserved regions) and LOF in CDS not encoding PFAM (Data S1). Additionally, we considered evidence for active transcription, quantified using aligned RNA-seq data from ten organs (NCBI BioProject PRJNA517635 ^5^) (see “*Gene model prediction, annotation, and curation”* section above). Gene counts were estimated with STAR v2.7.11b ^88^ (“--quantMode GeneCounts” flag) and edgeR v4.2.2 ^125^ was used to compute counts per million (CPM). A gene was deemed expressed if CPM ≥ 0.1 in ≥1 of the ten organs. To integrate this data, gene models were parsed to flag genes with annotated coding sequence (CDS) and to derive merged CDS intervals. Then the genome coordinates of putative LOF variants were intersected with these intervals in R ^122^, classifying each site as falling in an expressed CDS, non-expressed CDS, or non-CDS region (one hit per site, with expressed-CDS taking precedence when multiple overlaps occurred). Subsequent analyses were then performed only with the most stringent expressed-CDS subset.

As a proxy of individual genetic load, variants derived in Alpine ibex (“ibex-load”) and variants derived in domestic goat (“goat-load”) were defined as follows (see also Figure 4A). For both species, only variants fixed ancestral (either reference or alternative allele) in all individuals of the dataset, excluding domestic goat, hybrids, and Alpine ibex, were considered. For “ibex-load,” a site was required to be fixed ancestral among all domestic goats, and for “goat-load,” a site was required to be fixed ancestral among all Alpine ibex. All frequencies and filtering steps were performed using VCFtools v0.1.16 ^116^.

Next, for all LOF sites within expressed CDS, the number of heterozygotes and homozygotes derived (including both if the reference or the alternative allele was defined as derived, see above) were summed up in PLINK v2.0 ^117^. The number of heterozygotes represented the masked LOF load and the sum of heterozygotes and 2*homozygotes derived represented the total load ^51^. Absolute counts were used to quantify introgression of LOFs. For species comparisons, we accounted for species differences in genetic diversity by calculating a standardized LOF by putatively neutral variation. For this standardization, the same sums were taken for all sites annotated as modifier (putatively neutral) within expressed CDS. Standardized LOF stats were then defined as the ratio LOF stats / modifier stats, and a Welch two-sample t-test was performed to compare standardized LOF load between domestic goat and Alpine ibex.

### Copy number variations

The evaluation of copy number variations (CNVs) on the MHC complex was conducted on the aligned sequences using the software CNVkit version 0.9.10 ^126^. The mean read depth was calculated for all samples using the function *batch* and using the Alpine ibex sample as reference, with the calculation performed in intervals of approximately 1 Kbp. The weighted interval average across all samples belonging to the same species was evaluated using the function *reference*. Finally, a graphical representation of the copy number status at MHC loci was produced using the function *heatmap*.

### Phylogenetic analyses

Haplotype sequences of the DRA and DRB genes were obtained by extracting individual genomic sequences from the phased variant VCF file (see section “*Chromosome painting*”) using the consensus function in bedtools v2.30.0 ^87^ and the R package GenomicFeatures v1.58.0 ^127^. Multiple sequence alignments were generated with MUSCLE v3.8 ^128^ on the EMBL-EBI web server, using default parameters. Phylogenetic trees were imported, annotated and visualized with the R package ggtree ^129^ v3.14.0.

### Selection scan

The whole-genome VCF produced in the section “*SNP genotyping”* was prefiltered allowing for 5% missingness. It was then phased and imputed for missing genotypes with Beagle v27Feb25.75f ^121^. From the phased data, Alpine ibex, domestic goat and hybrid subsets were extracted, and haplotype-based selection scans were performed using selscan v2.1.1 ^130^, focusing on nSL ^131^ (within-population haplotype decay), because the statistic is robust to the absence of a recombination map and well suited for detecting incomplete selective sweeps in recently bottlenecked or admixed populations ^130^. While nSL is sensitive to gene flow, it is considered to be more robust than other statistics ^131^. Normalized |nSL| scores were computed for each group and local enrichment of extreme |nSL| values (top 1%) was quantified in sliding windows (200 kb, 5 kb step) to identify regions with an excess of unusually long haplotypes. This window-based enrichment was then compared across groups, with a particular focus on chromosome 23, where the MHC region is located.

### Simulation of demographic history, hybridization, and mutation load

The forward-in-time, age-structured individual-based simulator Nemo-age v0.32.0 ^132^ was used to model the consequences of hybridization between domestic goats and a bottlenecked Alpine ibex population. Simulations consisted of two patches (populations): a small, severely bottlenecked ibex population (pop1) and a large, stable goat population (pop2). Age structure and survival/fecundity was modelled using a seven-class life table reflecting Alpine ibex-like life history ^133,134^, with age-specific survival and fecundity encoded in a transition matrix and polygynous mating (mating_system = 2). Population regulation was imposed by patch-specific carrying capacities: pop1 was initially large (K = 1000), underwent a severe bottleneck to N ≈ 80 and later recovered to K = 500, whereas pop2 remained large (K = 5000).

Genomes comprised 1,624 biallelic loci subject to selection against deleterious mutations (mutation rate 5×10⁻ ⁵ and back mutation rate 5×10^−7^), with multiplicative fitness effects. These loci were randomly distributed across 29 *in silico* chromosomes as in the reference genome, each 200cM in length. Following Grossen et al. ^5^, the selection coefficients of 1,450 putatively deleterious loci were drawn from a gamma distribution with a mean of 0.01 and a shape parameter of 0.3 resulting in s < 1% for 99.2% of all loci ^135^. A negative relationship between s and hs was assumed ^136^ using the exponential equation h = exp(−51*s)/2 with a mean h set to 0.37 following Gilbert et al. ^137^. To include segregating high-impact alleles, 174 additional large-effect LOF mutations were added to the above-described set: 87 with s=1 and 87 with s=0.5, hs was set to 0 (strongly deleterious, fully recessive). Initial allele frequencies were set to 0.0014, following Grossen et al. ^5^. Neutral genetic diversity was tracked using 522 biallelic neutral loci (on average 18 per chromosome, mutation rate 5×10⁻ ⁵), from which expected heterozygosity was calculated at predefined generations.

To investigate the effect of different hybridization scenarios, dispersal was varied from no dispersal, to 1% or 2.5% of pop2 dispersing to pop1. In other words, these simulated dispersal rates refer to the percent of individuals moving per generation from the source population (domestic goat) to the target (Alpine ibex). The high dispersal rate scenarios reflect the findings that one of the two hybrid swarms might have been formed following the abandonment of an entire goat herd in the same region ^34^.

Furthermore, dispersal either occurred as a pulse, only during a short, time-limited gene flow pulse after the bottleneck, for 4 generations while set to zero before and after, or dispersal was kept at a constant rate of 1% for 180 generations (until end of simulations). Each set of simulations was run for 10,200 generations, where a burn-in of 10’000 generations allowed new mutations to occur. The bottleneck of pop1 occurred from generation 10,000 to 10,010, and gene flow began at generation 10,021 and either went until 10,024 (pulse) or 10,200 (end of the simulations, ongoing gene flow, as an extreme scenario).

To assess the consequences of maladaptive introgression disrupting local adaptation, we extended the simulations by incorporating local adaptation as a quantitative trait. Specifically, a single quantitative trait controlled by five loci was simulated, with Gaussian stabilizing selection acting toward population-specific optima (set to −2 for ibex and +2 for goats). Trait evolution followed a mutation–selection framework with a per-locus mutation rate of 1×10⁻⁴ and mutation variance of 0.05. Distinct local optima were defined for the two populations, such that Alpine ibex (pop1) were adapted to their local environment, whereas gene flow from goats (pop2) introduced alleles potentially maladapted to this optimum. Selection strength was governed by a Gaussian fitness function with variance parameter ω² = 30. The environmental deviation variance of the trait phenotype was set to 0 (default), such that phenotype was fully determined by genotype. This maladaptive introgression scenario was implemented to the 1% pulse dispersal regime (1% dispersal for four generations), allowing exploration of the effects of maladaptive allele introgression.

For each replicate, Nemo-age ^132^ recorded per-generation population size, population differentiation, average fitness per patch, neutral allele frequencies per locus, and the full genotype-based load (sum of deleterious effects) for both populations. A total of 50 independent replicates were run, summarizing genetic load measures and neutral heterozygosity trajectories in pop1 and pop2 across time (see also Data S3).

The full Nemo-age ^132^ parameter files are also available on GitHub [https://github.com/fgcilingir/ibex_hybrid_load].

As done for the empirical data, simulated genetic load was estimated as derived allele counts in homozygotes (realized load), heterozygotes (masked load) and the sum of both (total load) ^51^. Severe genetic load (as a proxy of LOFs) was defined as all mutations with s>0.1. Standardization was performed using putatively neutral mutations with s<0.0001.

