## Supplementary material for "Hybridization load introduced by Alpine ibex hybrid swarms": Document S1

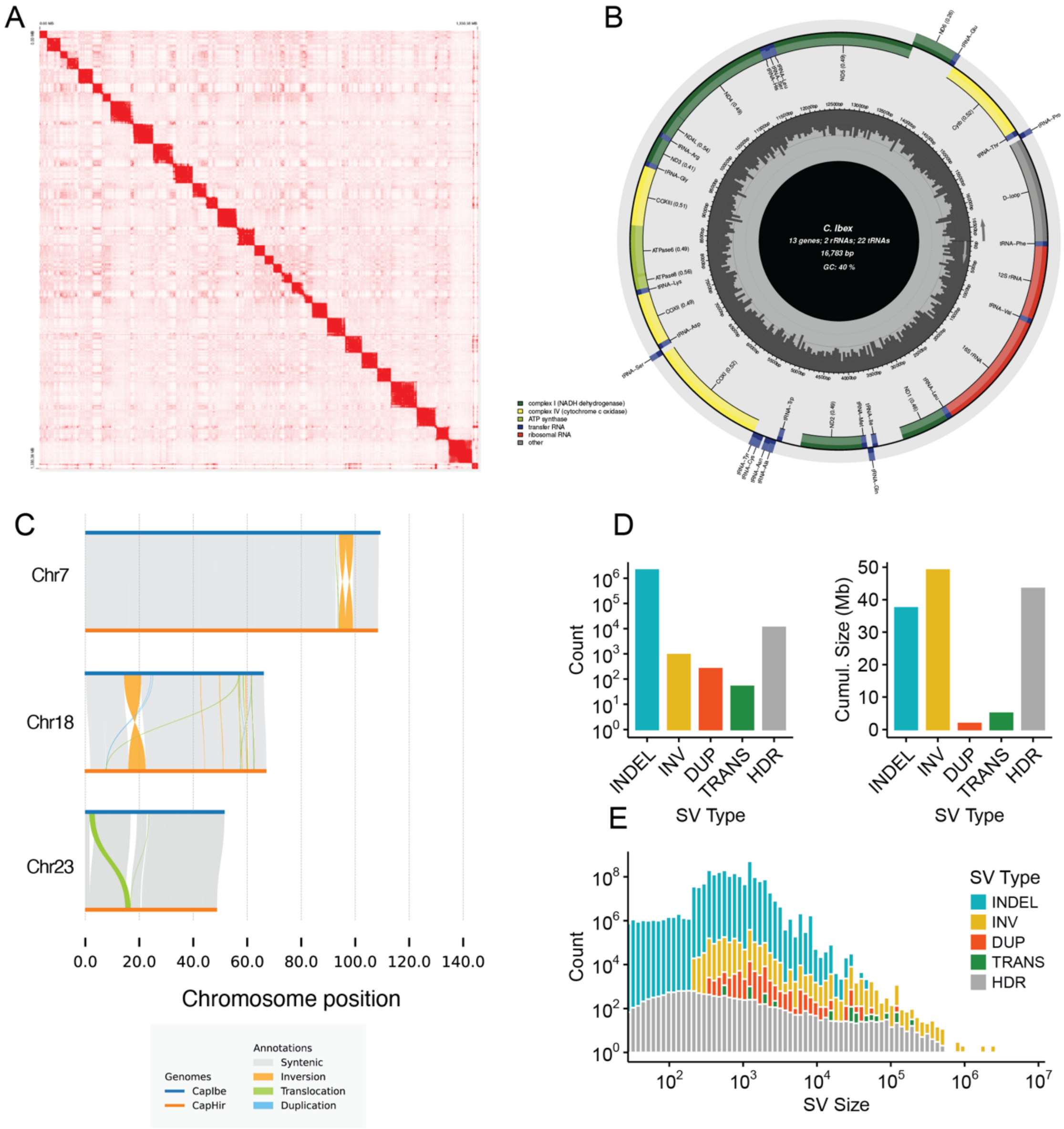


**Figure S1 A)** HiC contact map showing the data used for genome scaffolding. **B)** Schema of the Alpine ibex mitochondrial genome. **C)** Major translocations between domestic goat and Alpine ibex observed on chromosomes 7, 18, and 23 based on SyRI**. D)** Total counts and cumulative sizes of the structural variants identified by SyRI between the Alpine ibex and domestic goat genomes. **E)** Size distribution of translocations shown as a histogram from the same variants as in D (INDEL: insertion/deletion, INV: inversion, DUP: duplication, TRANS: translocation, HDR: highly divergent region). Related to Figure 1, STAR Methods

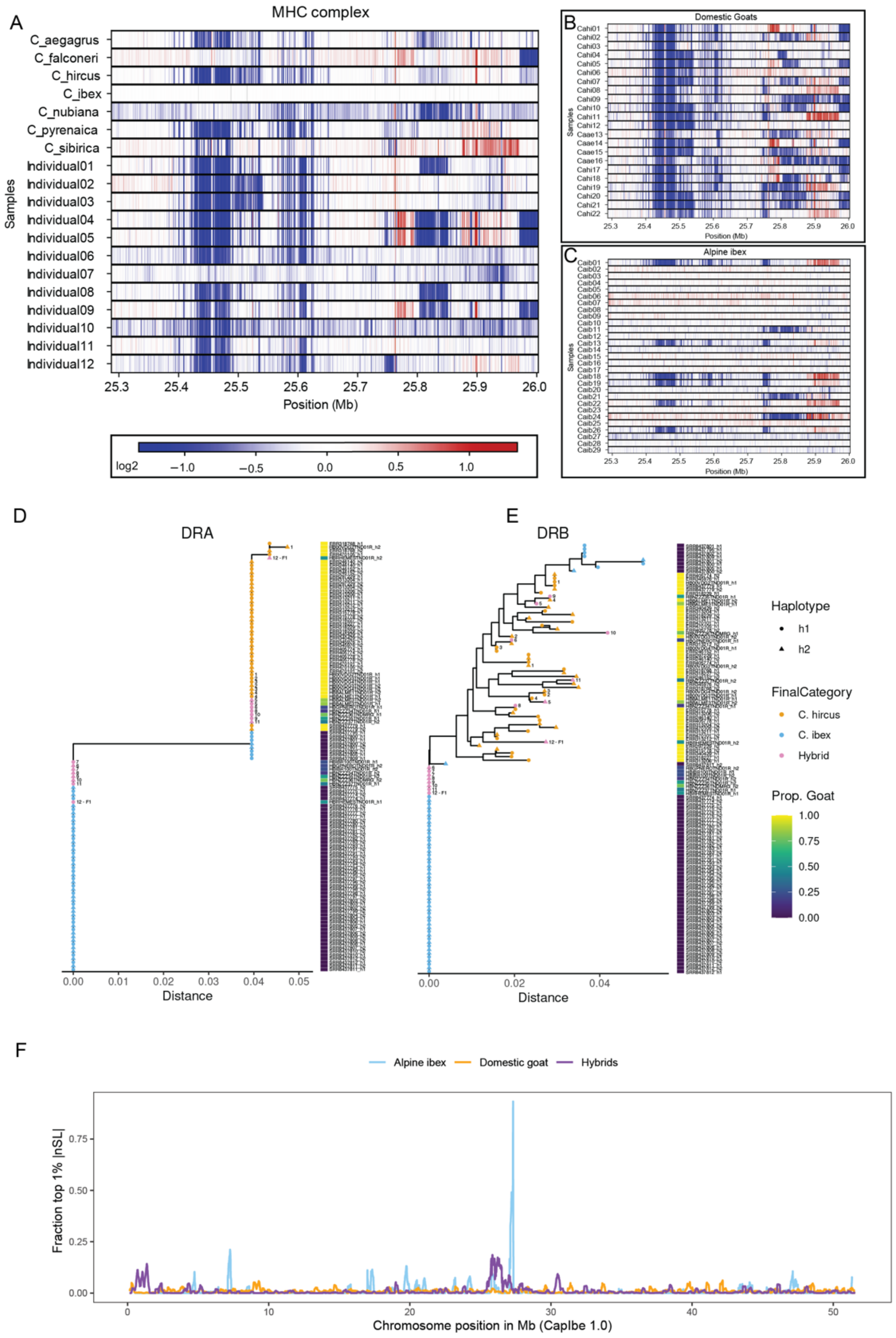


**Figure S2 (A–C)** Copy-number (CN) variation across the MHC class II locus. Heatmaps display normalized read-depth estimates across the MHC class II region (blue = deletions; red = duplications). **A)** CN profiles for wild Capra species (*C. aegagrus*, *C. falconeri*, *C. nubiana*, *C. pyrenaica,* and *C. sibirica*) together with the domestic goat and newly sequenced hybrid and ibex individuals. A double deletion at 25.4–25.5 Mb is present in most species but absent from *C. ibex* and *C. falconeri*. Additional variation, including localized deletions and duplications, occurs along the remainder of the MHC class II region. **B)** Domestic goats consistently carry the 25.4–25.5 Mb double deletion and exhibit further CN heterogeneity across the locus. **C)** Alpine ibex show markedly reduced CN variability, with largely uniform copy number across the region. A subset of individuals (Caib01,

Caib13, Caib18, Caib19, Caib22, Caib26) carries the goat-like double-deletion haplotype, consistent with historical introgression at immune loci. **(D-E)** Phylogenetic trees obtained from the alignments of the DRA **D)** and DRB **E)** proteins from *C. hircus* (orange), *C. ibex* (light blue) and hybrid individuals (purple). The heatmap shows the proportion of goat genome, as calculated with the chromosome painting analysis. **F)** Sliding-window enrichment of extreme |nSL| genome-wide shown with the fraction of SNPs exceeding the genome-wide 99th percentile of normalized |nSL| in 200 Kb windows (5 Kb step) across chromosome 23 for Alpine ibex (light blue), domestic goats (orange), and hybrids (purple). All groups exhibit low background levels of high |nSL| values across most of the scaffold, but Alpine ibex and hybrids show a pronounced, localized enrichment peak within the MHC region. Related to Figure 2.

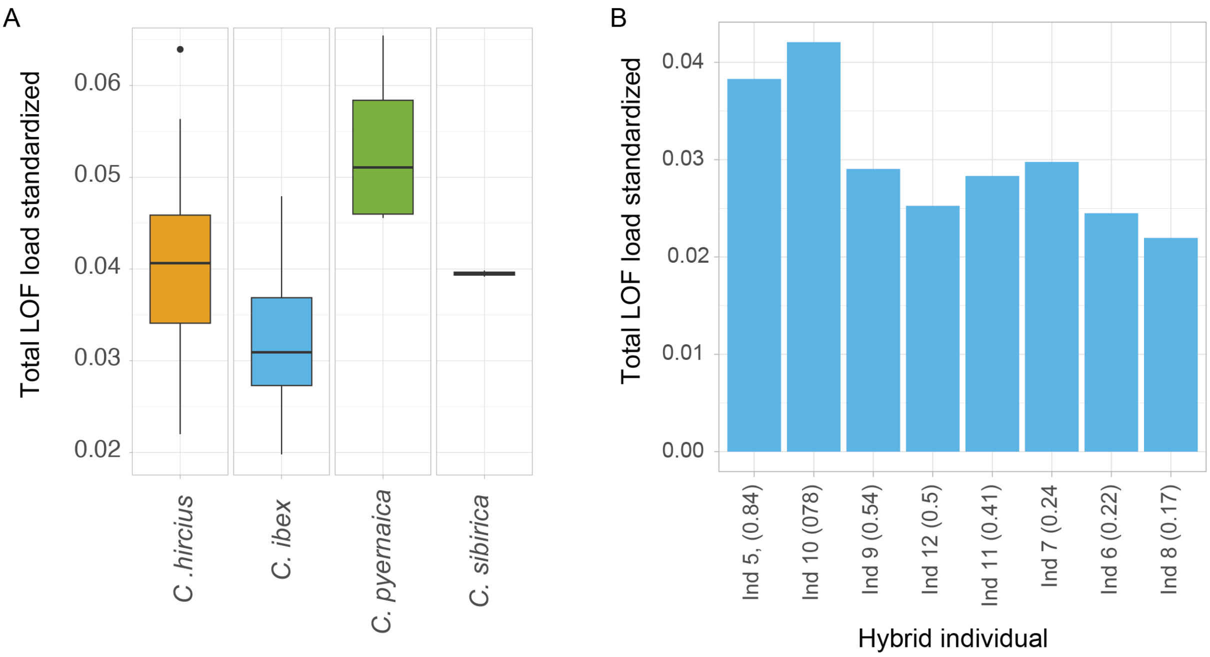


**Figure S3** A) Standardized total individual counts of loss-of-function alleles (standardized by the total counts of modifier alleles) for domestic goats (*C. hircus*), Alpine ibex (*C. ibex*), Iberian ibex (*C. prenaica*) and Siberian ibex (*C. sibirica*). B) Standardized total counts of loss-of-function alleles for each hybrid individual ordered by proportion of domestic goat (indicated in parentheses). Individuals with proportion domestic goat lower than 0.5 are back-crosses into Alpine ibex, which show no reduction of loss-of-function load. Related to Figure 4.
