## Supplementary material for "Hybridization load introduced by Alpine ibex hybrid swarms": Data S2

Read depth profiles of already available whole genome sequencing samples of Alpine ibex (N=29), domestic goat (N=18), wild goat (N=4), Iberian ibex (N=4), and 12 newly sequenced individual samples.

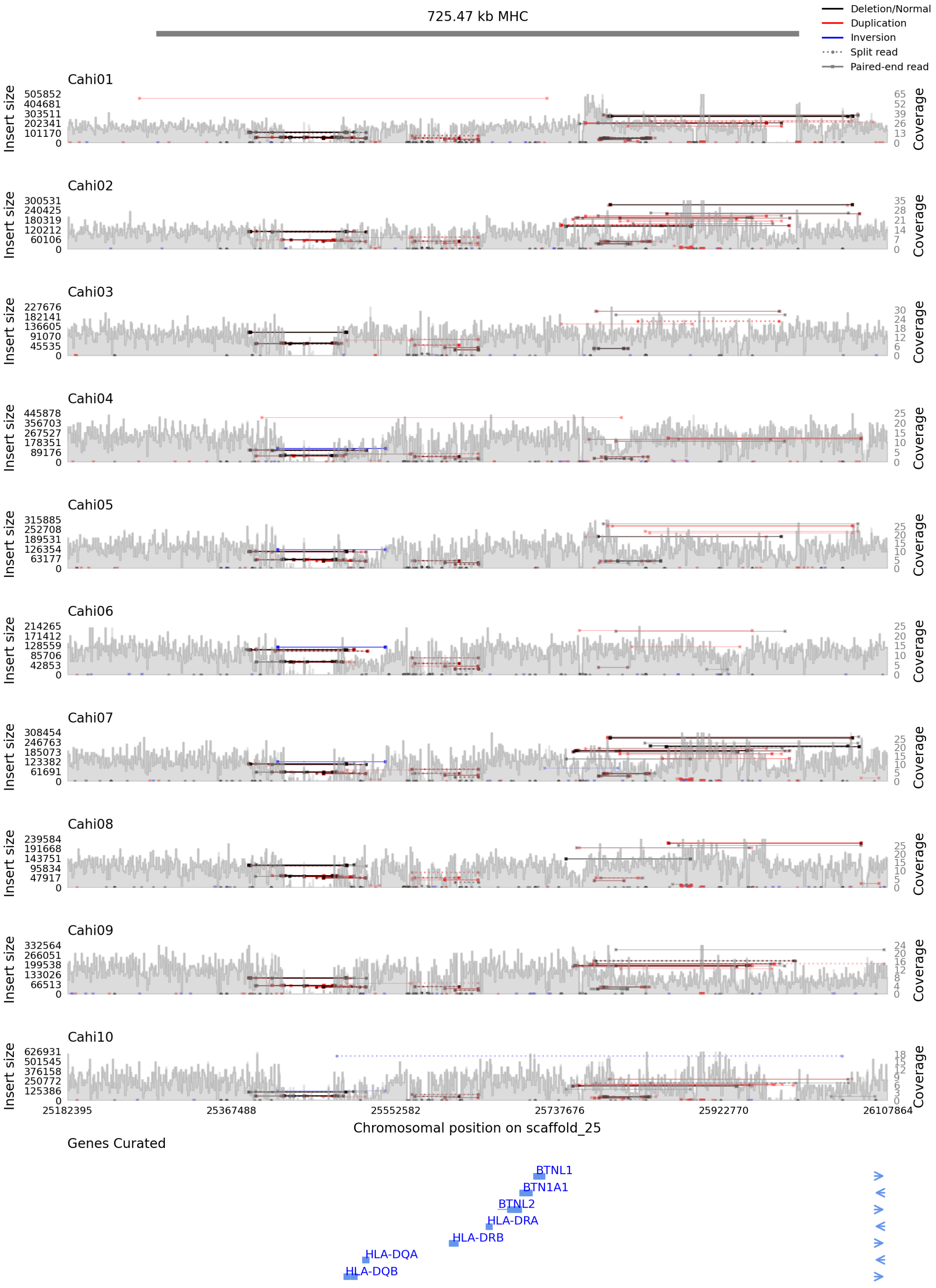

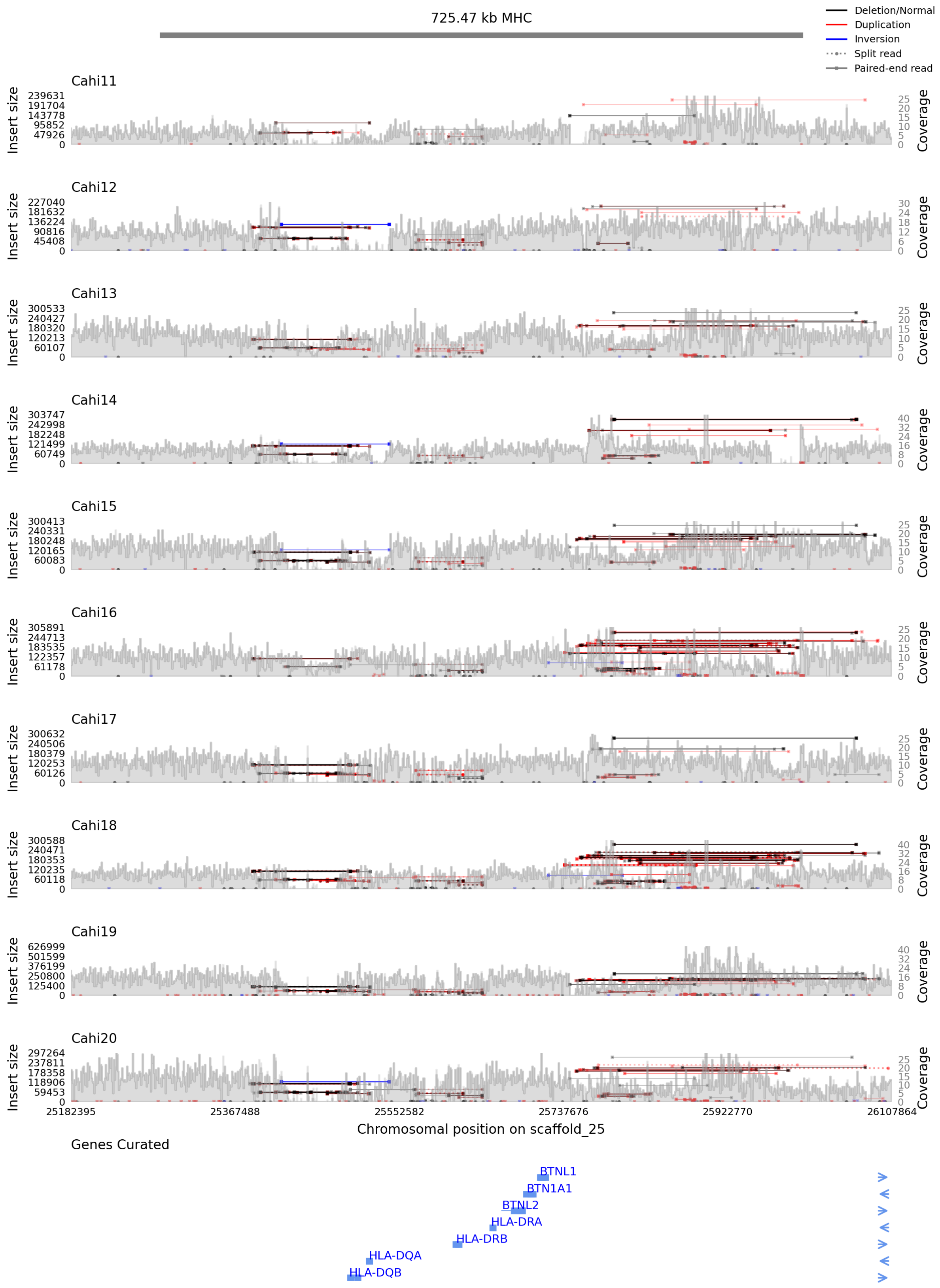

725.47 kb MHC
Deletion/Normal
Duplication
Inversion
Split read
Paired-end read
Cahi11
Cahi12
Cahi13
Cahi14
Cahi15
Cahi16
Cahi17
Cahi18
Cahi19
Cahi20
Insert size
Coverage
25182395
25367488
25552582
25737676
25922770
26107864
Chromosomal position on scaffold_25
Genes Curated
BTNL1
BTN1A1
BTNL2
HLA-DRA
HLA-DRB
HLA-DQA
HLA-DQB

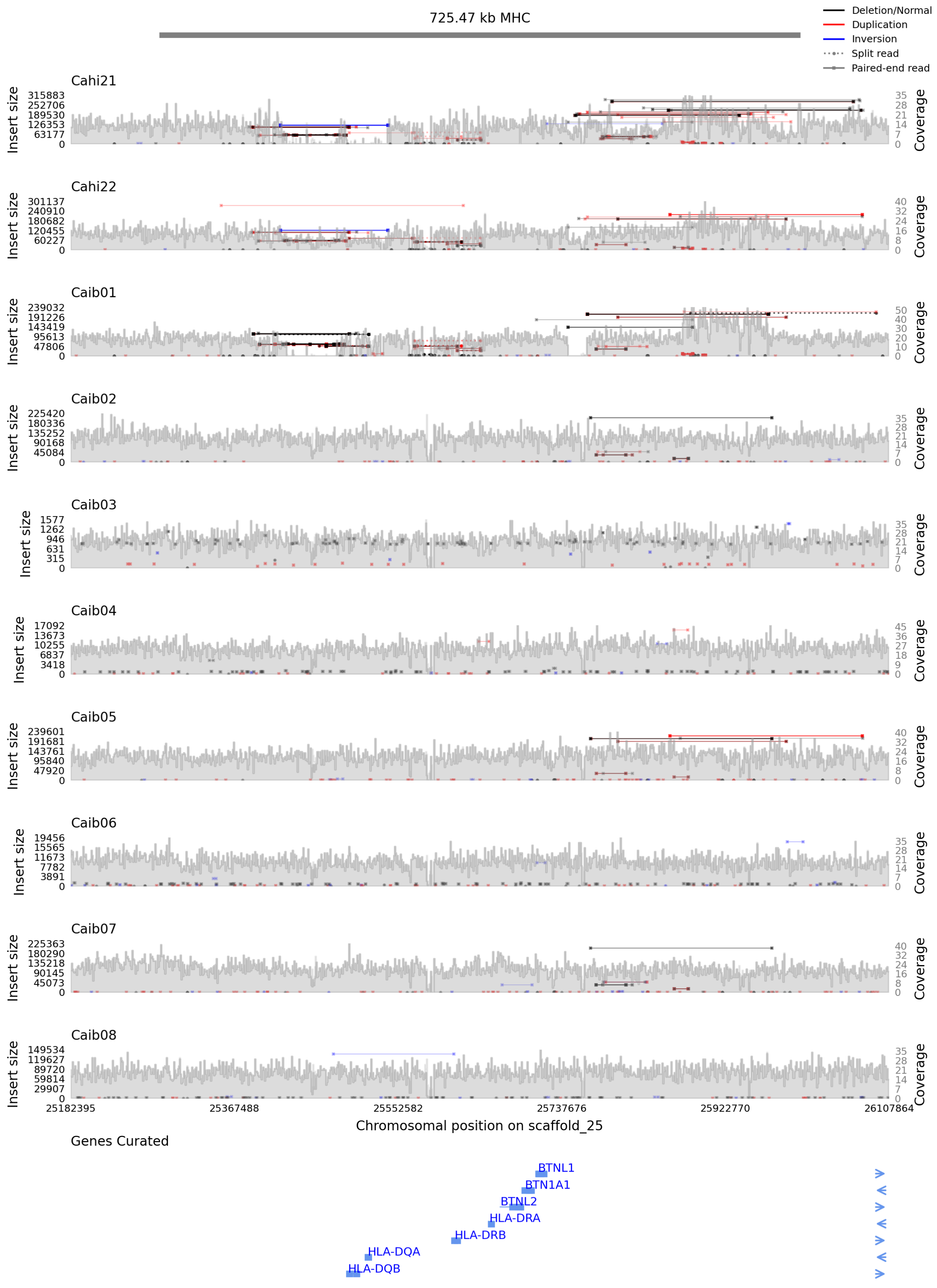
725.47 kb MHC
Deletion/Normal
Duplication
Inversion
Split read
Paired-end read
Cahi21
Cahi22
Caib01
Caib02
Caib03
Caib04
Caib05
Caib06
Caib07
Caib08
Insert size
Coverage
25182395
25367488
25552582
25737676
25922770
26107864
Chromosomal position on scaffold_25
Genes Curated
BTNL1
BTN1A1
BTNL2
HLA-DRA
HLA-DRB
HLA-DQA
HLA-DQB

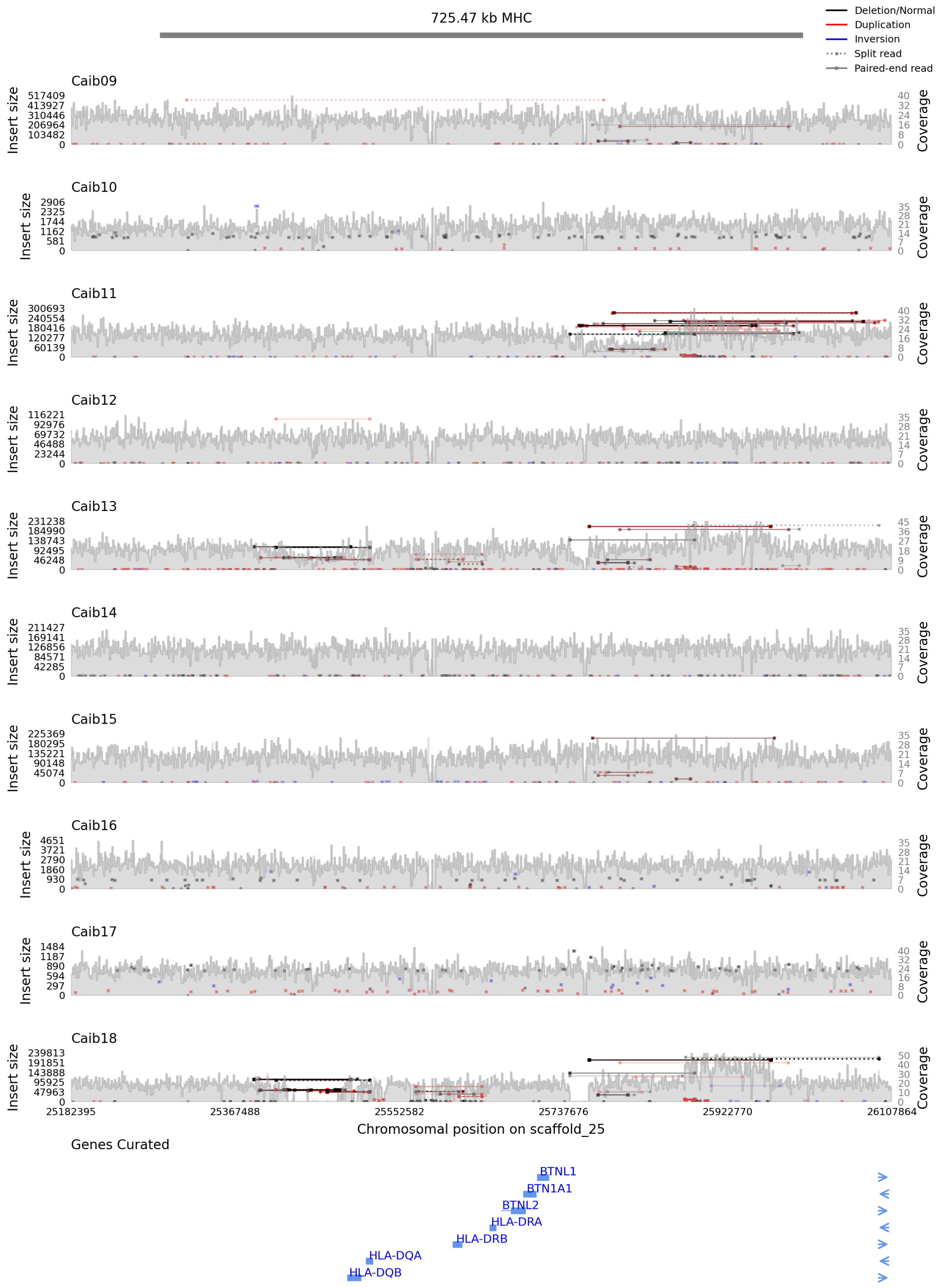

725.47 kb MHC
Deletion/Normal
Duplication
Inversion
Split read
Paired-end read
Caib09
Caib10
Caib11
Caib12
Caib13
Caib14
Caib15
Caib16
Caib17
Caib18
Insert size
Coverage
25182395
25367488
25552582
25737676
25922770
26107864
Chromosomal position on scaffold_25
Genes Curated
BTNL1
BTN1A1
BTNL2
HLA-DRA
HLA-DRB
HLA-DQA
HLA-DQB

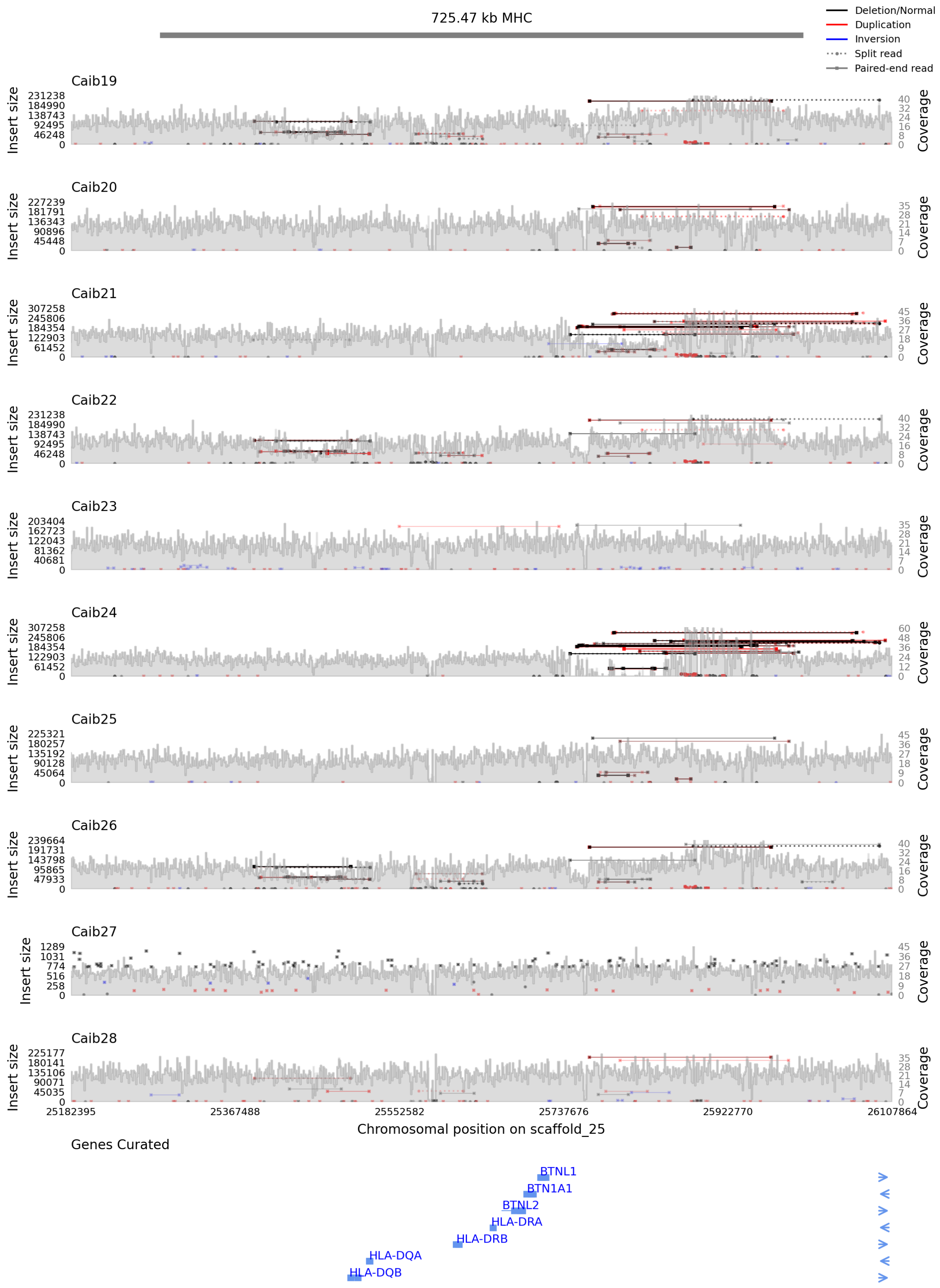
725.47 kb MHC
Deletion/Normal
Duplication
Inversion
Split read
Paired-end read
Caib19
Caib20
Caib21
Caib22
Caib23
Caib24
Caib25
Caib26
Caib27
Caib28
Insert size
Coverage
25182395
25367488
25552582
25737676
25922770
26107864
Chromosomal position on scaffold_25
Genes Curated
BTNL1
BTN1A1
BTNL2
HLA-DRA
HLA-DRB
HLA-DQA
HLA-DQB

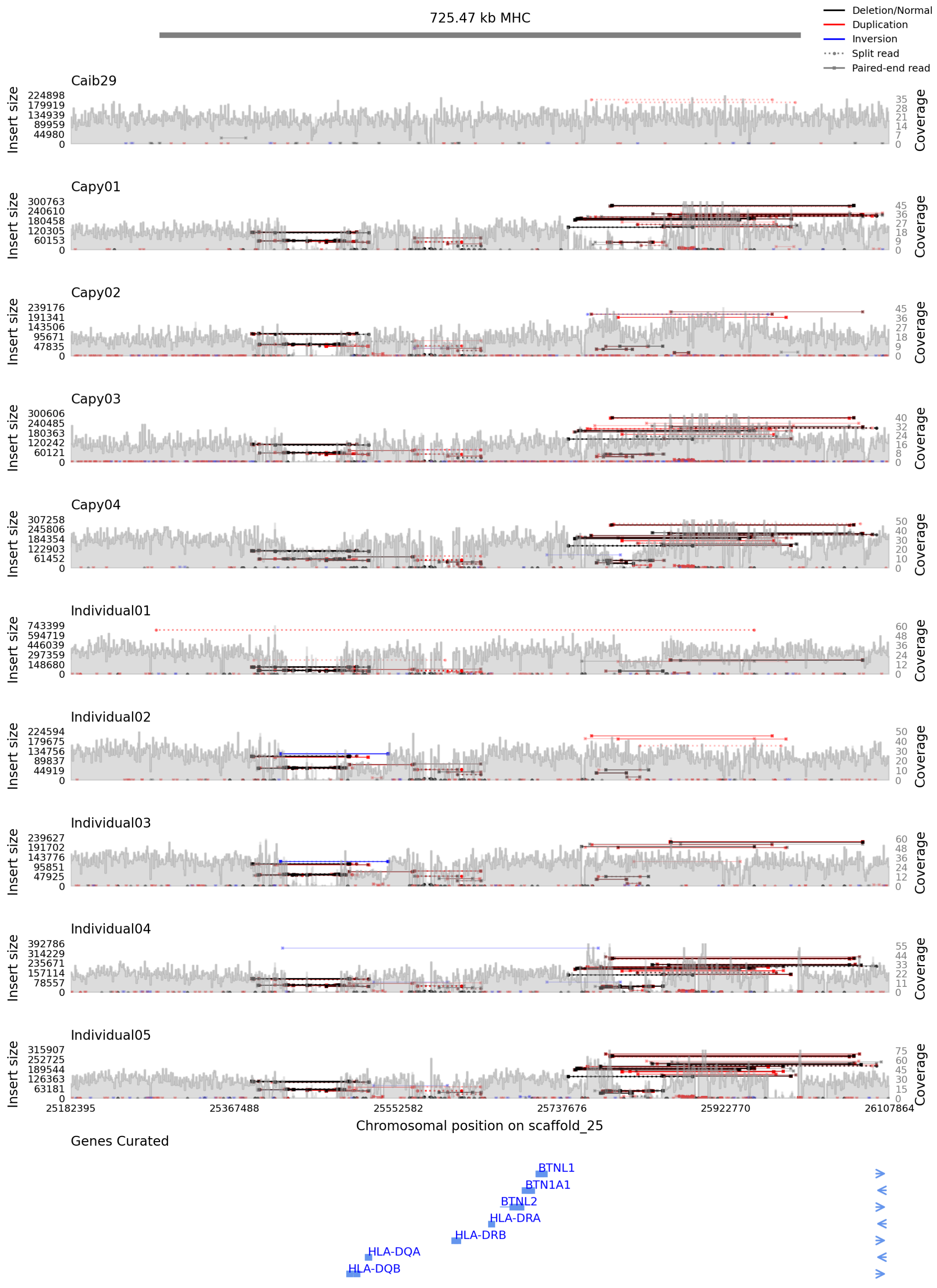
725.47 kb MHC
Deletion/Normal
Duplication
Inversion
Split read
Paired-end read
Caib29
Capy01
Capy02
Capy03
Capy04
Individual01
Individual02
Individual03
Individual04
Individual05
Insert size
Coverage
25182395
25367488
25552582
25737676
25922770
26107864
Chromosomal position on scaffold_25
Genes Curated
BTNL1
BTN1A1
BTNL2
HLA-DRA
HLA-DRB
HLA-DQA
HLA-DQB

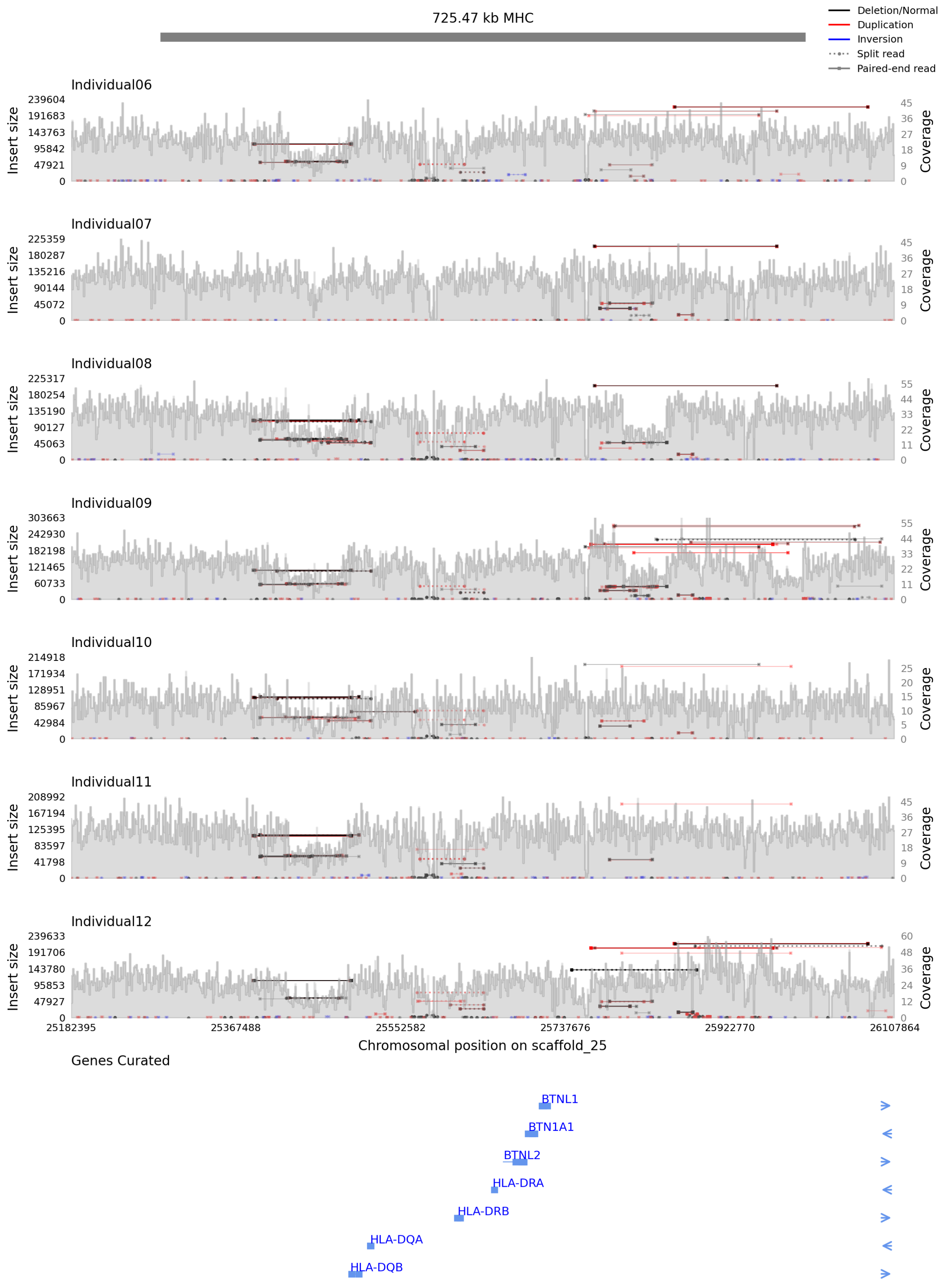
725.47 kb MHC
Deletion/Normal
Duplication
Inversion
Split read
Paired-end read
Individual06
Insert size
239604
191683
143763
95842
47921
0
Coverage
45
36
27
18
9
0
Individual07
225359
180287
135216
90144
45072
0
45
36
27
18
9
0
Individual08
225317
180254
135190
90127
45063
0
55
44
33
22
11
0
Individual09
303663
242930
182198
121465
60733
0
55
44
33
22
11
0
Individual10
214918
171934
128951
85967
42984
0
25
20
15
10
5
0
Individual11
208992
167194
125395
83597
41798
0
45
36
27
18
9
0
Individual12
239633
191706
143780
95853
47927
0
60
48
36
24
12
0
25182395
25367488
25552582
25737676
25922770
26107864
Chromosomal position on scaffold_25
Genes Curated
BTNL1
BTN1A1
BTNL2
HLA-DRA
HLA-DRB
HLA-DQA
HLA-DQB
