## Supplementary material for "Hybridization load introduced by Alpine ibex hybrid swarms": Data S3

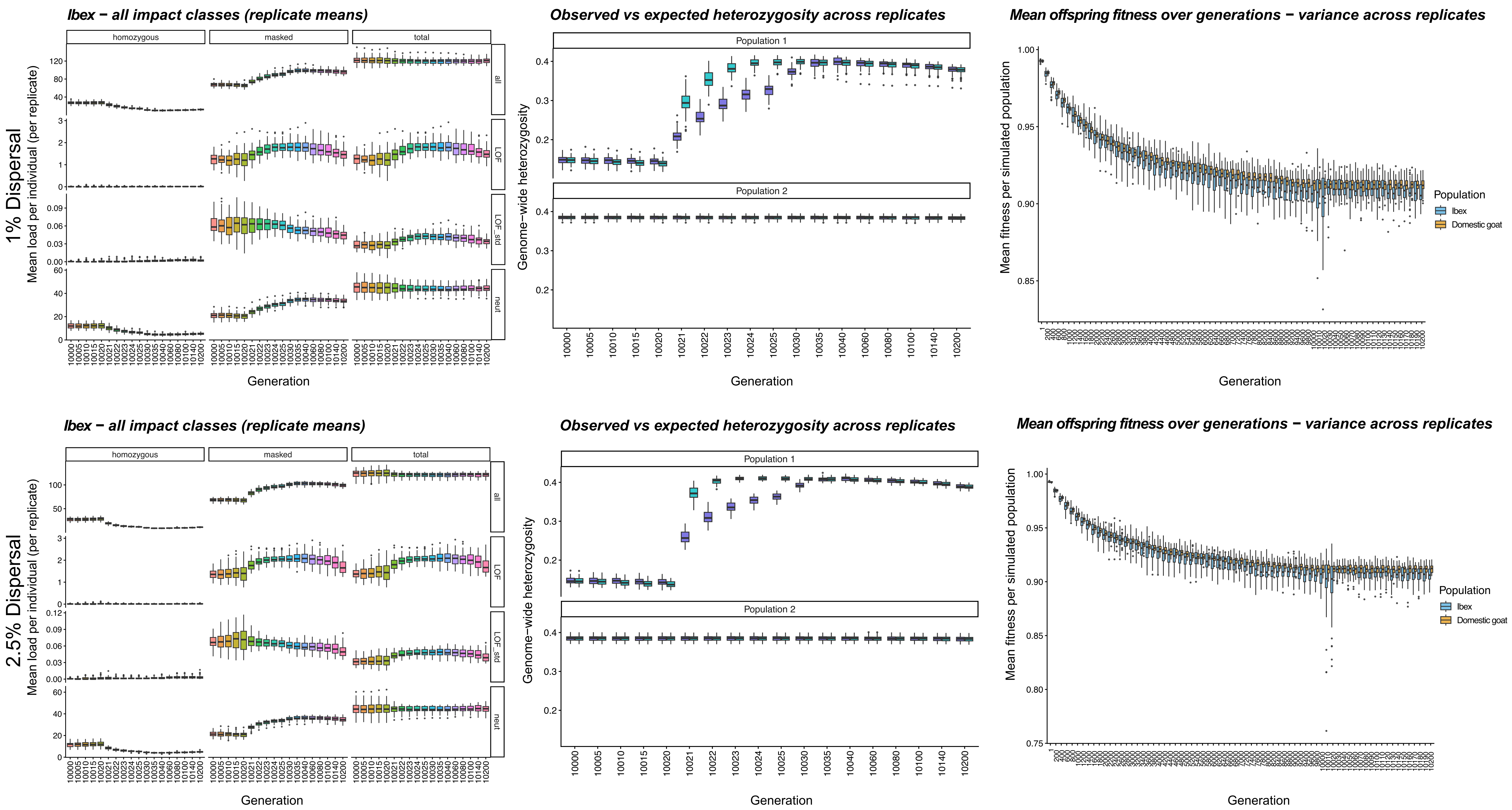
1% Dispersal
2.5% Dispersal
Ibex − all impact classes (replicate means)
homozygous
masked
total
all
LOF
LOF_std
neut
Mean load per individual (per replicate)
Generation
Observed vs expected heterozygosity across replicates
Population 1
Population 2
Genome−wide heterozygosity
Mean offspring fitness over generations − variance across replicates
Mean fitness per simulated population
Population
Ibex
Domestic goat

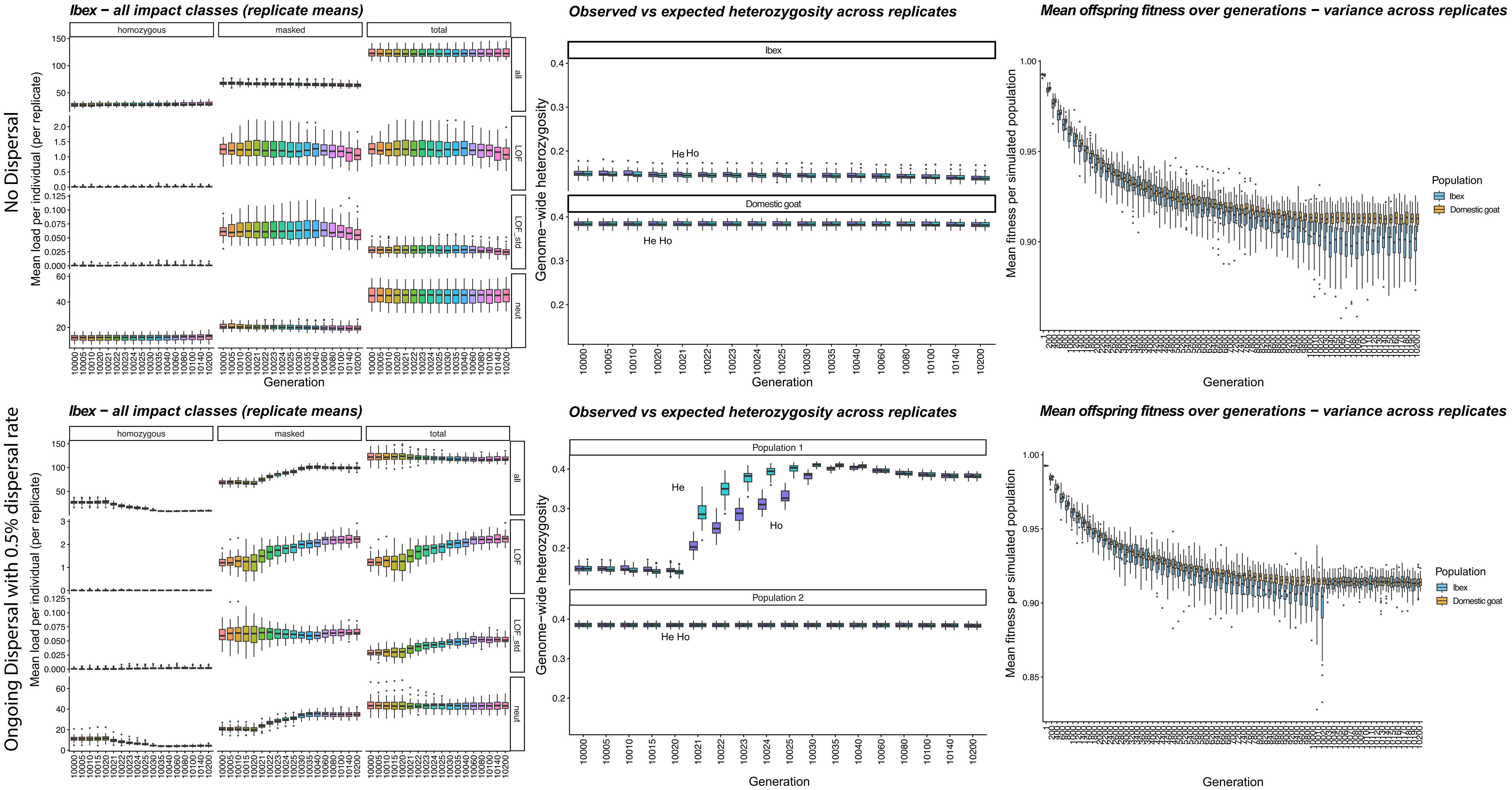
No Dispersal
Ongoing Dispersal with 0.5% dispersal rate
Ibex − all impact classes (replicate means)
homozygous
masked
total
all
LOF
LOF_std
neut
Mean load per individual (per replicate)
Generation
Observed vs expected heterozygosity across replicates
Ibex
Domestic goat
Population 1
Population 2
He
Ho
Genome−wide heterozygosity
Mean offspring fitness over generations − variance across replicates
Mean fitness per simulated population
Population
Ibex
Domestic goat

### Simulations also including adaptive loci

**Ibex − all impact classes (replicate means)**

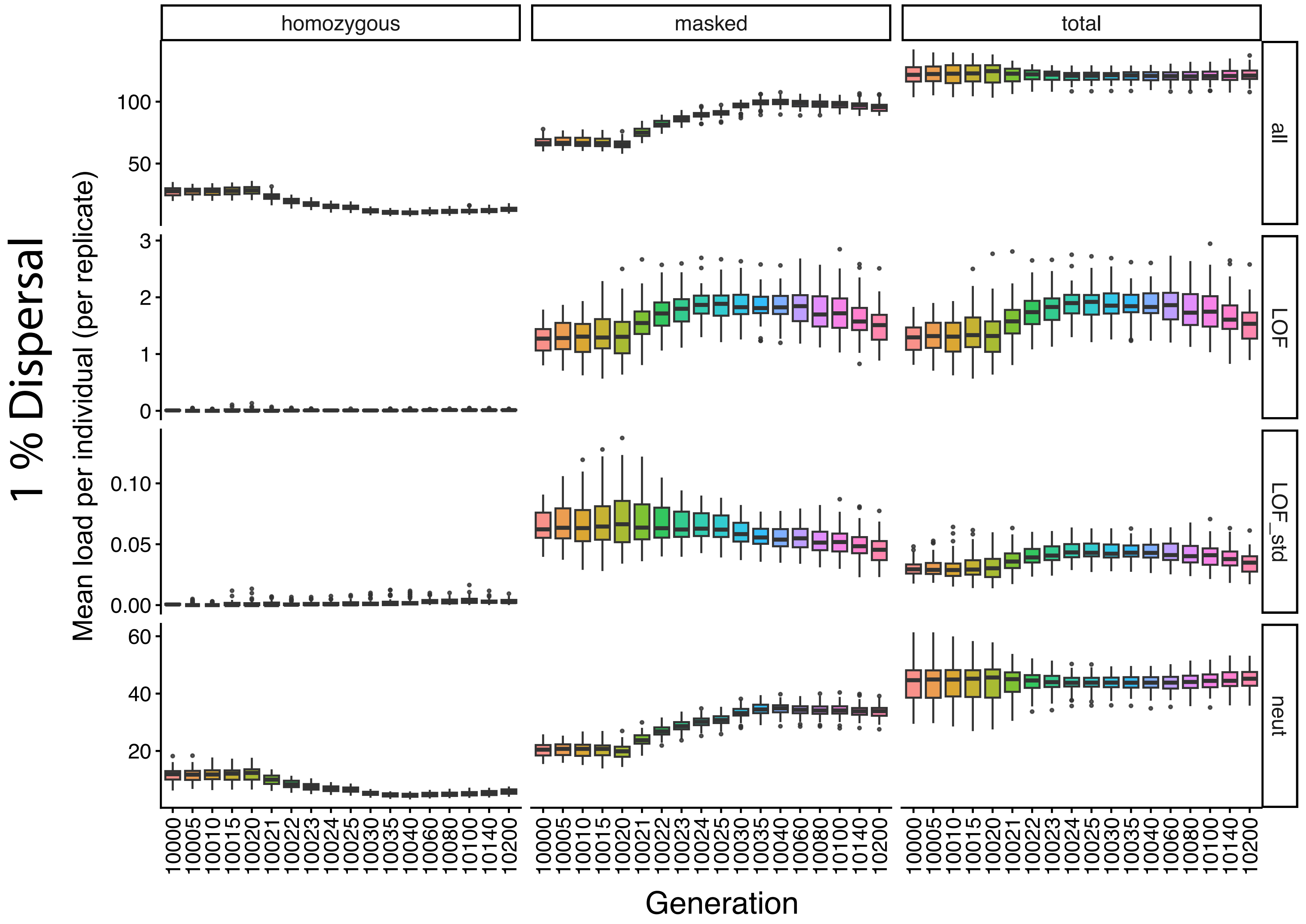


**Observed vs expected heterozygosity across replicates**

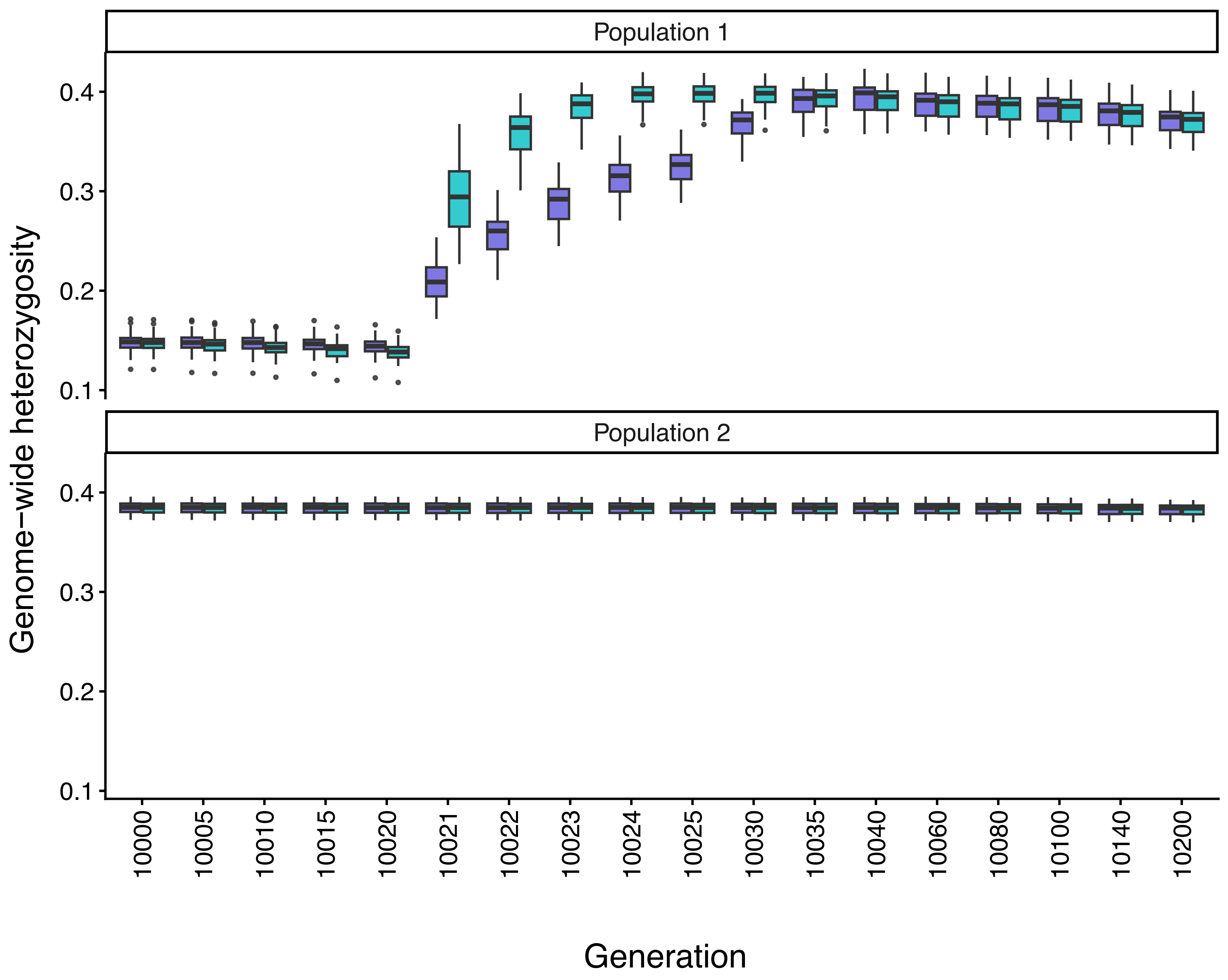


**Mean offspring fitness over generations − variance across replicates**

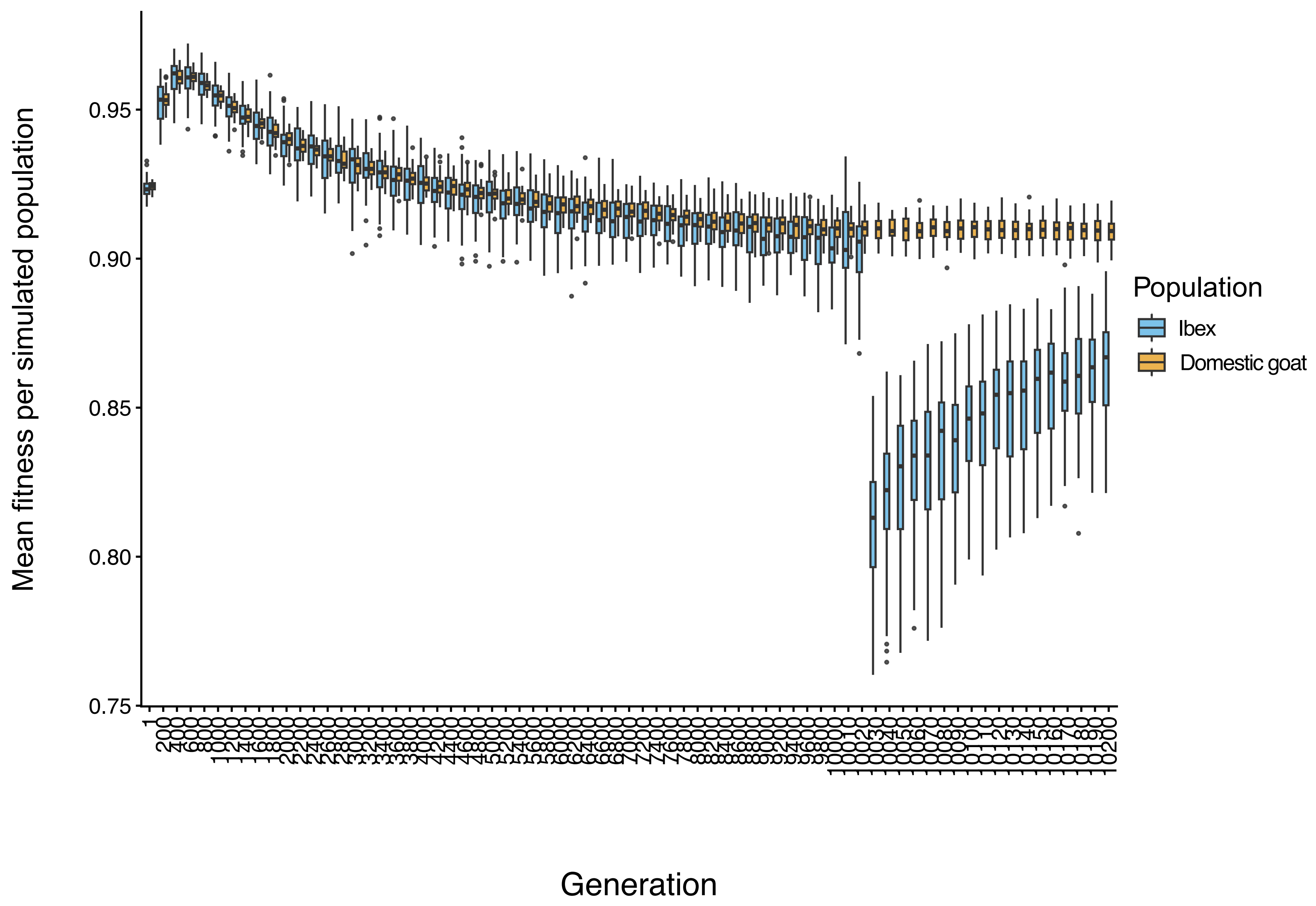
